# Inheritance of old mitochondria controls early CD8^+^ T cell fate commitment and is regulated by autophagy

**DOI:** 10.1101/2024.01.29.577412

**Authors:** Mariana Borsa, Ana Victoria Lechuga-Vieco, Amir H. Kayvanjoo, Yavuz Yazicioglu, Ewoud B. Compeer, Felix C. Richter, Hien Bui, Michael L. Dustin, Pekka Katajisto, Anna Katharina Simon

## Abstract

T cell immunity is impaired during ageing, particularly in memory responses needed for efficient vaccination. Autophagy and asymmetric cell division (ACD) are cell biological mechanisms key to memory formation, which undergo a decline upon ageing. However, despite the fundamental importance of these processes in cellular function, the link between ACD and *in vivo* fate decisions has remained highly correlative in T cells and in the field of mammalian ACD overall. Here we provide robust causal evidence linking ACD to *in vivo* T cell fate decisions and our data are consistent with the concept that initiation of asymmetric T cell fates is regulated by autophagy. Analysing the proteome of first-daughter CD8^+^ T cells following TCR-triggered activation, we reveal that mitochondrial proteins rely on autophagy for their asymmetric inheritance and that damaged mitochondria are polarized upon first division. These results led us to evaluate whether mitochondria were asymmetrically inherited and to functionally address their impact on T cell fate. For this we used a novel mouse model that allows sequential tagging of mitochondria in mother and daughter cells, enabling their isolation and subsequent *in vivo* analysis of CD8^+^ T cell progenies based on pre-mitotic cell cargo. Autophagy-deficient CD8^+^ T cells showed impaired clearance and symmetric inheritance of old mitochondria, suggesting that degradation events promote asymmetry and are needed to generate T cells devoid of old organelles. Daughter cells inheriting old mitochondria are more glycolytic and upon adoptive transfer show reduced memory potential, whereas daughter cells that have not inherited old mitochondria from the mother cell are long-lived and expand upon cognate-antigen challenge. Proteomic and single-cell transcriptomic analysis of cells inheriting aged mitochondria suggest that their early fate divergence relies on one carbon metabolism as a consequence of poor mitochondrial quality and function. These findings increase our understanding of how T cell diversity is early-imprinted during division and will help foster the development of strategies to modulate T cell function.

The MitoSnap model allows tracking of pre-mitotic and post-mitotic cell cargoes.
Both segregation and degradation (autophagy) contribute to the asymmetric inheritance of old mitochondria.
Old mitochondria impact cell metabolism and function.
Cells devoid of old mitochondria exhibit better memory potential *in vivo*.

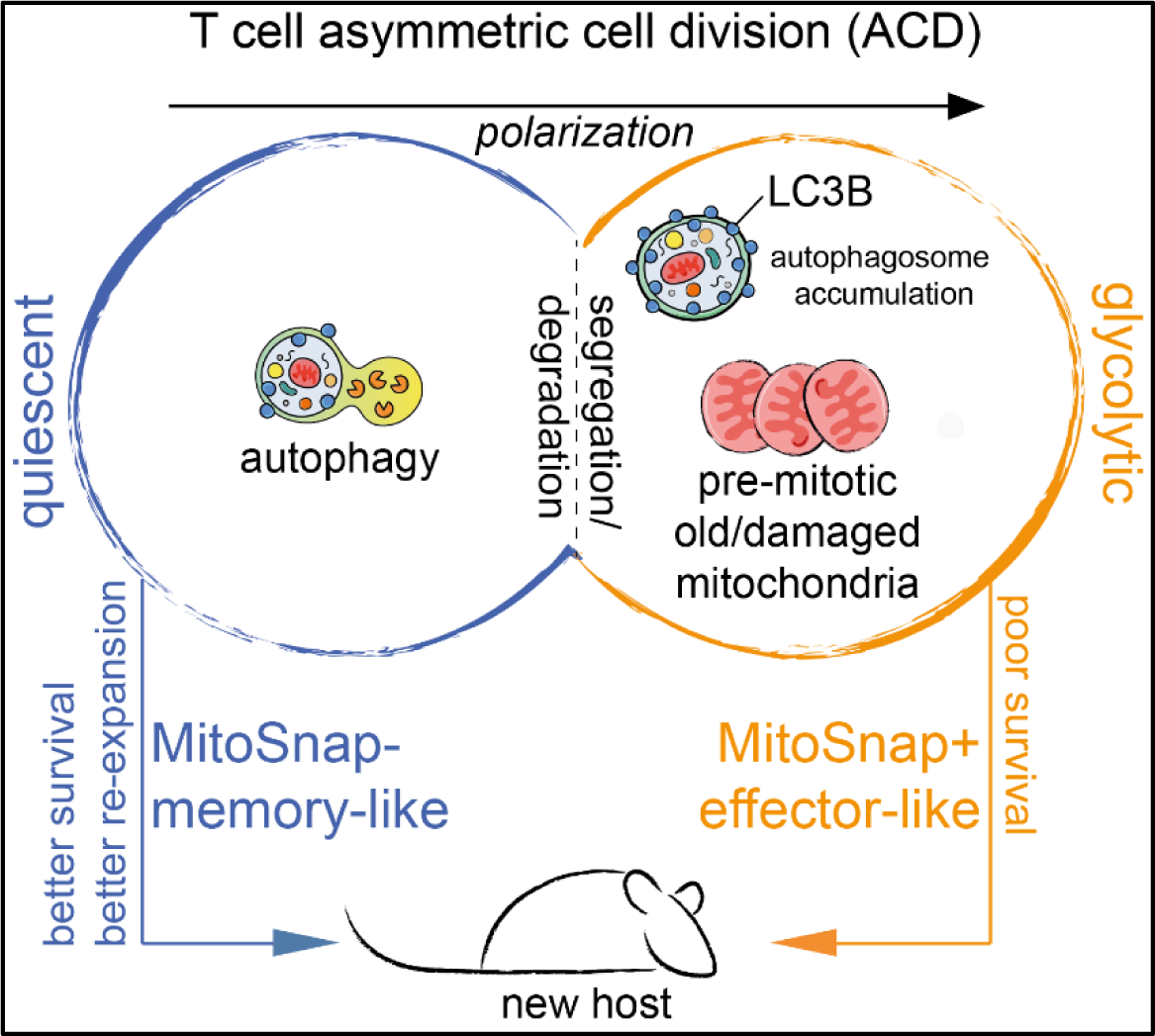

## Introduction

Efficient immune responses rely on the coordinated function of different immune cell types, which also requires generation of diversity within the same cell type. In the context of CD8^+^ T cells, one single cell is able to differentiate and generate progeny with heterogeneous fates upon activation^1,2^. Activation of a naïve T cell by its cognate antigenic epitope leads to the differentiation of short-lived effector cells that exert cytotoxic effector functions, and long-lived memory cells that self-renew and differentiate upon antigenic re-challenge and are central to vaccination efficacy. Despite increased understanding of mechanisms that contribute to fate decision during CD8^+^ T cell differentiation, there is still no consensus on when these decisions are made, and particularly how long-lived memory T cells are formed^3–5^.. Moreover, during aging T cell memory is severely impaired^6,7^, and senescent CD8^+^ T cell subsets that exhibit DNA damage, cell cycle arrest, mitochondrial dysfunction due to defective mitophagy^8^, and global poor effector function accumulate^9–11^. Amongst the cellular processes that benefit the formation and maintenance of memory CD8^+^ T cells but which are negatively impacted by ageing^6,12,13^, there are two highly conserved mechanisms: macroautophagy (hereafter termed autophagy) and asymmetric cell division (ACD).

Autophagy involves the recycling and degradation of cellular cargoes, which occurs via the engulfment of cellular components by double-membrane structures called autophagosomes, and their delivery to lysosomes for degradation. The regulation by autophagy of immune cell fate decision is cell- and context-dependent^14^. In CD8^+^ T cell differentiation, autophagy loss results in an impaired memory response^6,15,16^, which is at least partly caused by accumulation of damaged organelles^17,18^.

ACD has been well characterised in model organisms such as yeast, *Drosophila melanogaster* and *Caenorhabditis elegans*^19^, but evidence of the impact of this mechanism on cell fate in mammalian cells remains correlative and inconclusive^20^. In cells from the haematopoietic lineage, this is a consequence of technical limitations as *in vivo* functional readouts of sibling cells have relied on cell cargoes that do not directly influence fate decisions and/or show dynamic and variable expression. Thus, a critical question remains: is inherited material synthesized post-cell division, or is it inherited asymmetrically? Here we address this question using CD8^+^ T cells. ACD in CD8^+^ T cells is important for the generation of two distinct cell types, through the early generation of effector-like and memory-like daughter cells^21^ that occurs from the first mitosis after naïve T cell activation by high-affinity TCR stimulation^22,23^. The daughter cells emerging from ACD inherit several layers of asymmetry, including the differential expression of surface markers, transcription factors, divergent metabolic activity and translation^24–27^. However, a direct link between asymmetric inheritance of pre-mitotic T cell cargo and the future fate of emerging daughter cells *in vivo* has not been made. Thus, solid causal evidence linking ACD to fate decisions is lacking in T cells and in the field of mammalian ACD in general.

Because it is unclear whether cell cargo degradation can contribute to cell division asymmetries, we performed an integrated functional analysis of the contribution of autophagy and ACD to CD8^+^ T cell differentiation. We identified damaged mitochondria as asymmetrically inherited cargo, and this asymmetry is further deepened on mitophagy. Then, using a novel mouse model that allows specific labelling of mitochondria before and after cell division, we tracked mitochondrial inheritance and biogenesis, ensuring that this cell cargo was not perturbed by post-mitotic changes in CD8^+^ T cell progenies. This novel tool allowed us to follow the presence of pre-mitotic mitochondria by imaging and flow cytometry, and evaluate the impact of mitochondrial inheritance by proteomics, scRNAseq and *in vivo* transfer of daughter cells. Our results suggest that autophagy contributes to the generation of early divergent cell fates by promoting both clearance and asymmetric partitioning of old mitochondria. Furthermore, we are the first to unequivocally draw a causal link between the inheritance of cell cargo to future fate commitment, as old mitochondria caused poor memory potential in CD8^+^ T cell immune responses. Our findings offer new insight into how T cell diversity is imprinted early during cell division, and how organelle ageing regulates CD8^+^ T cell metabolism and function, facilitating more refined therapeutic approaches to T cell modulation.

## Results

### Divergent proteome and mitochondrial inheritance in CD8^+^ T cell mitosis relies on autophagy

Asymmetric cell division in CD8^+^ T cells results in the unequal inheritance of different cell cargoes that culminates in divergent transcriptomes between daughter cells^27–30^. We aimed to broaden our understanding of early events of asymmetric segregation by assessing the global proteome of first-daughter CD8^+^ T cells. To that end, we used CD8 as a surrogate marker to classify effector-like (CD8^hi^) and memory-like (CD8^lo^) progenies. Briefly, we isolated naïve CD8^+^ T cells from spleens and lymph nodes of wild-type (WT) C57BL/6 mice, labelled them with a cell trace dye and activated these cells for 36-40 h on anti-CD3, anti-CD28 and Fc-ICAM-1 coated wells. First-daughter CD8^+^ T cells were sorted into CD8^hi^ or CD8^lo^ populations as previously described^28^ (Fig.1A). Cells were washed and pellets were used for quantitative label-free high-resolution mass spectrometry. >6000 proteins were identified and the proteomic ruler method was used to calculate both protein mass and copy numbers of each protein per cell^31^. We did not observe differences in total protein mass between CD8^hi^ and CD8^lo^ daughter cells (Fig. S1A). However, we identified several proteins that were enriched in one of these two populations, as represented by fold-change in protein copy number between effector-like and memory-like cells following first mitosis (Fig. 1B). Amongst the top 50 identified targets in each group, we found several proteins with a role linked to cell metabolism, mitochondrial function and biogenesis, which are highlighted in bold. Because mitochondria are known to contribute to T cell fate^32,33^, we decided to focus on these organelles. We aimed to validate the results obtained from this unbiased approach by imaging mitochondria in mitotic cells and emerging siblings. By electron microscopy, we could neither observe any differences in mitochondrial content (Fig. 1C), nor any differences in mitochondrial architecture. The inheritance of mitochondria by daughter cells during mitosis has been superficially investigated with conflicting results^25,26,34^. Thus, we evaluated whether mitochondrial fitness is different between CD8^+^ T cell siblings. Using the cell permeable probe MitoSOX, we imaged mitochondria producing high levels of reactive oxygen species (ROS, a readout of damaged organelles), and observed that CD8^hi^ (effector-like) daughter cells had a higher abundance of mitochondrial ROS production (Fig. 1D). Because damaged mitochondria are targets of autophagy - a mechanism known to benefit memory CD8^+^ T cells - and known to undergo an age-related decline, we interrogated whether mitophagy contributes to this unequal distribution and quantified MitoSOX inheritance in autophagy-deficient CD8^+^ T cells. Using non-inducible autophagy-deleted CD8^+^ T cells *(Atg7*^fl/fl^ *Cd4*^Cre^), we observed that the immune synapse (IS) area and TCR clustering were distinct between the autophagy-sufficient and -deficient CD8^+^ T cells (Fig. S1B). As it has been described that IS formation and TCR-affinity and signalling strength are crucial for asymmetric T cell division^21–23^, we excluded that any differences in T cell activation due to loss of Atg7 interferes with ACD readouts by using an inducible model of autophagy deletion (*Atg16l1*^fl/fl^ *Ert2*^Cre^). Here activation happens with functional autophagy, as *Atg16l1* is deleted only upon *in vitro* Z-4-Hydroxytamoxifen (4OHT) treatment (Fig. S1C), and Cre-recombination events do not result in immediate ATG16L1 loss. We analyzed mitotic CD8^+^ T cells by confocal microscopy at 36-40 h post-activation, and found that autophagy loss abolishes the asymmetric inheritance of damaged (MitoSOX+) mitochondria (Fig. 1D). To evaluate whether the autophagic machinery itself is polarized during cell division, we evaluated the expression of the autophagy-marker LC3B. LC3B is the lipidated and membrane-bound version of Microtubule-associated protein 1A/1B-light chain 3 (LC3), which functions in autophagy substrate selection and autophagosome biogenesis and is a target of degradation itself during the autophagic process when no lysosomal inhibitor is added^14^. As observed for MitoSOX, LC3B was co-inherited by CD8^hi^ (effector-like) daughter cells, suggesting that this daughter cell performs less autophagy/mitophagy, which leads to the accumulation of autophagy targets. To confirm this, we also evaluated the inheritance of LC3B in CD8^+^ T cells from aged mice, known to show poor ACD potential and low autophagy levels^12,15,16^. We found that ageing leads to the symmetric inheritance of LC3B (Fig. 1E). Finally, these results could be correlated with the proteome of CD8^hi^ and CD8^lo^ daughter cells generated from autophagy-deficient (inducible *Atg16l1* deletion) and aged cells (Fig. 1F), since: i) the numbers of differentially inherited proteins were lower than the ones observed in WT cells (Fig 1B), and ii) we found fewer and different proteins linked to mitochondrial function amongst the differentially-inherited in autophagy-deficient and aged CD8^+^ T cells. Together these findings highlight the relevance of autophagy in the establishment of asymmetric inheritance patterns. Interestingly, the pool of enriched proteins found in CD8^hi^ and CD8^lo^ daughter cells was very small in old mice and no mitochondrial proteins were found, perhaps because both ACD and autophagy decline with age.

**Figure 1.**
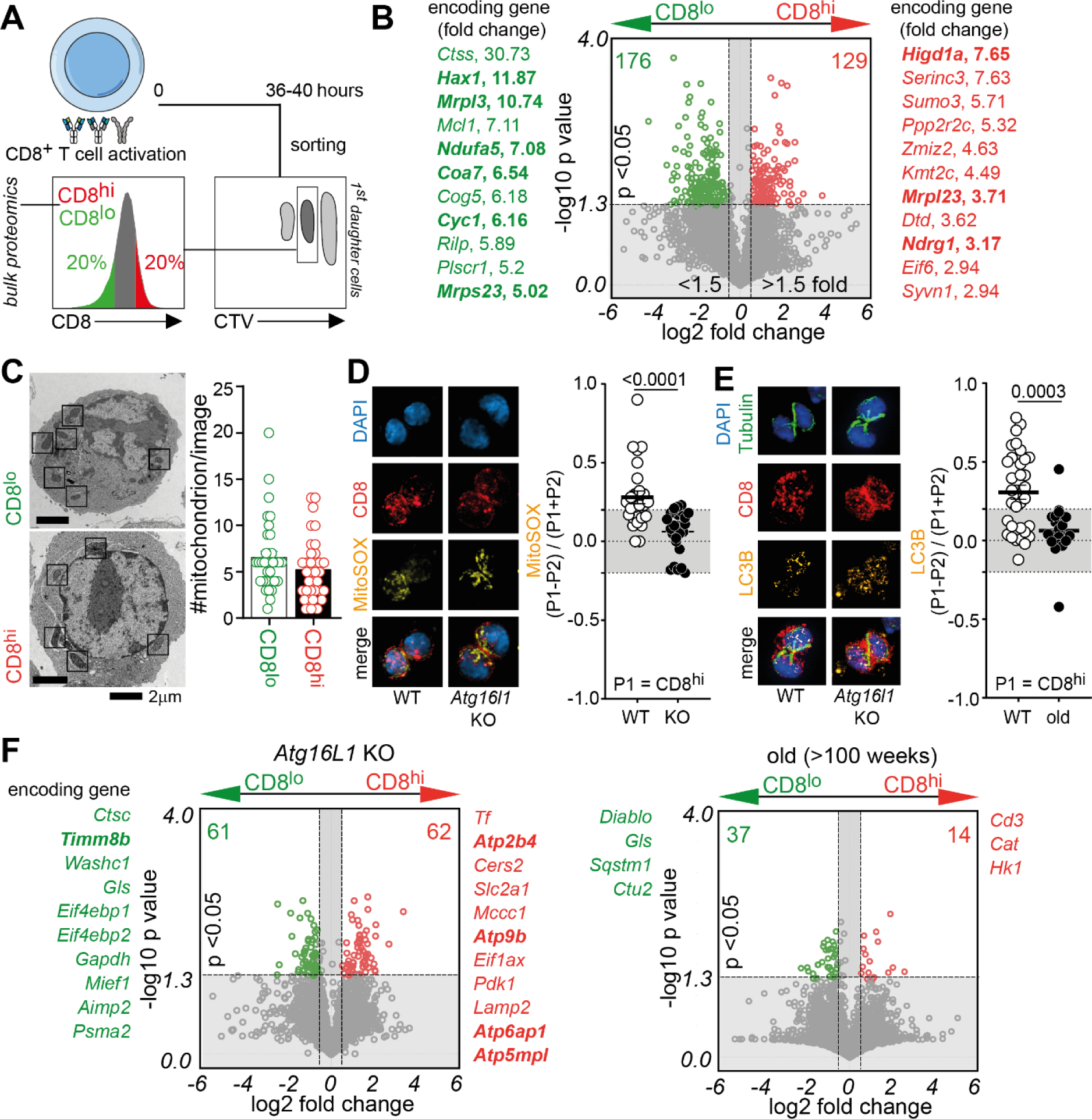
Autophagy regulates asymmetries in CD8^+^ T cell mitosis. **(A)** Experimental layout: CTV-labelled naïve CD8+ T cells were activated on anti-CD3, anti-CD28 and Fc-ICAM-1 coated plates for 36-40 h. Cells were harvested and stained with anti-CD8 antibodies. First-daughter cells were identified as the first peak of CTV dilution (in reference to undivided cells). CD8^hi^ and CD8^lo^ cells were sorted as populations expressing 20% highest or lowest CD8, respectively, as previously described^28^. Cells pellets were frozen and stored until being processed for proteomics analysis. **(B)** Volcano plot showing differentially inherited proteins by CD8^hi^ and CD8^lo^ daughter-cells. Data pooled from 4 samples done in 2 independent experiments. Each sample had cells originally harvested from 2-3 mice. Encoding genes for proteins amongst the top 50 differentially expressed in CD8^lo^ and CD8^hi^ daughter cells are highlighted in green and red, respectively. Genes in bold have their function linked to mitochondrial metabolism and function. **(C)** Representative transmission electron microscopy images from CD8^hi^ and CD8^lo^ daughter cells emerging from the first mitosis following naïve CD8^+^ T cell activation (left). Number of mitochondria per image/slice was calculated (right). **(D)** Representative images of WT (*Atg16l1*^fl/-^ *Ert2*^Cre^) and autophagy KO (*Atg16l1*^fl/fl^ *Ert2*^Cre^) mitotic CD8^+^ T cells 36-40 h post-activation. Autophagy depletion was achieved by culturing cells in presence of 500 nM Z-4-Hydroxytamoxifen (4OHT). Inheritance of MitoSOX was calculated as previously described^28^. Any values above or below the grey area in the graph were considered asymmetric. Data are represented as mean ± SEM. Statistical analysis was performed using an unpaired two-tailed Student’s *t* test. Exact P values are depicted in the figure. **(E)** Representative images of mitotic CD8^+^ T cells from young (8-16 weeks-old) and old (>100 weeks-old) mice 36-40 h post-activation. Inheritance of LC3B, a marker of autophagosomes, was calculated in each group. Data are represented as mean ± SEM. Statistical analysis was performed using an unpaired two-tailed Student’s *t* test. Exact P values are depicted in the figure. **(F)** Volcano plot showing differentially inherited proteins by CD8^hi^ and CD8^lo^ daughter-cells from *Atg16l1*^fl/-^ *Ert2*^Cre^ (post-tamoxifen inducible depletion of autophagy) and old (>100 weeks) mice. Data pooled from 2-4 samples. Each sample had cells originally harvested from 2-3 mice. Encoding genes for proteins amongst the top 50 differentially expressed in CD8^lo^ and CD8^hi^ daughter cells are highlighted in green and red, respectively, for each type of sample. Genes are ordered from top to bottom in decreasing fold-change values. Genes in bold have their function linked to mitochondrial metabolism and function. Proteomics volcano plots were done using Tableau.

### Inheritance of old mitochondria is autophagy-dependent

To functionally address whether damaged organelles play a role as fate determinants during ACD *in vivo*, we took advantage of the MitoSnap murine model (MGI:6466976; *Omp25*-SnapTag^fl/-^ *Ert2*^Cre^), which allows mitochondria to be followed from the mother cell, hereafter named ‘old’ based on the permanent fluorescent labelling of a SnapSubstrate targeted to mitochondria via OMP25. SnapTag is a modified DNA repair enzyme that can covalently bind to different cell-permeable substrates linked to fluorophores. Sequential labelling of SnapTag expressing cells allows separation by flow cytometry of different populations based on patterns of organelle inheritance. We optimized the timelines to discriminate between older and younger organelles in CD8^+^ T cells. Briefly, naïve CD8^+^ T cells were isolated, activated overnight on anti-CD3, anti-CD28 and Fc-ICAM-1 coated plates in the presence of Z-4-Hydroxytamoxifen (4OHT) to induce SnapTag expression, and labelled with two different fluorescently labelled SnapSubstrates at 16 h (‘old’, before 1^st^ mitosis), and 36 h (‘young’, post-mitotic) post-stimulation. Incubation with an unlabelled SnapSubstrate (SnapBlock) was done immediately before the second labelling, guaranteeing that young organelle structures had emerged from recent biogenesis. Downstream analysis was done 2 h post-young labelling (Fig. 2A)^35,36^. With the MitoSnap system we can unequivocally link the inheritance of labelled (old) mitochondria to an event of asymmetric segregation of a cell cargo that was present >24 h before cell division. Furthermore, this cargo is not affected by recent transcriptional, translational or anabolic events of biogenesis. SnapTag labelled mitochondria co-localized with Tom20+ structures by fluorescence confocal microscopy, confirming the specificity of the SnapTag chemistry (Fig. 2B). Analysis of first-daughter CD8^+^ T cells by flow cytometry revealed the emergence of two main populations inheriting either both old and young mitochondria or exhibiting no SnapTag labelling (Fig. 2C). Importantly, SnapTag negative cells result from degradation or segregation of mitochondria, as labelling efficiency is close to 100% (Fig. S2A). To confirm whether these two populations result from segregation into the daughter cells or degradation, we generated autophagy-deficient MitoSnap mice (*Atg16l1*^fl/fl^ *Omp25*-SnapTag^fl/-^ *Ert2*^Cre^). We labelled old SnapTag mitochondria from autophagy-sufficient and autophagy-deficient CD8^+^ T cells and evaluated their loss over several cell divisions. Autophagy-deficient MitoSnap CD8^+^ T cells only lost the old mitochondria in 3.6% of all dividing cells, as opposed to 23% in WT conditions (Fig. 2D). To directly observe segregation events, mitotic MitoSnap CD8^+^ T cells were analyzed by fluorescence confocal microscopy, revealing that asymmetric segregation of old mitochondria occurs in WT but not KO cells and therefore relies on autophagy (Fig. 2E). Importantly, we confirmed that old mitochondria are MitoSOX+ (Fig. 2F). To further dissect the role of mitophagy in the generation of MitoSnap-cells, we sorted MitoSnap+ and MitoSnap-cells following first CD8^+^ T cell division and put them back in culture without any further TCR stimulation for 3 days in T cell medium containing IL-2, IL-7 and IL-15, which are cytokines that promote survival and memory maintenance (Fig. 2G). Using MitoSnap-conditions as reference negative controls, we observed that WT cells that were originally MitoSnap+ became MitoSnap-. However, most of the MitoSnap+ autophagy-deficient CD8^+^ T cells maintained their SnapTag labelling (Fig. 2H). Finally, aiming to comprehend the mitochondrial content of autophagy-sufficient and -deficient CD8^+^ T cells at the organelle level, we enriched mitochondrial fractions of both types of cells at 40 h post-activation (as in Fig. 2A) and performed flow cytometry analysis. We gated on mitochondria based on their size and Tom20 expression and observed that mitochondrial units are larger in *Atg16l1*-deficient cells (Fig. 2A, Fig. S2B). Supporting the role of mitophagy in the clearance of aged mitochondria, we observed that autophagy-deficient cells had a higher proportion of mitochondria preserving old organelle labelling in comparison to their WT counterparts (Fig. 2I). These results suggest that the emergence of MitoSnap-cells relies both on segregation and degradation events and that autophagy plays a role in both mechanisms.

**Figure 2.**
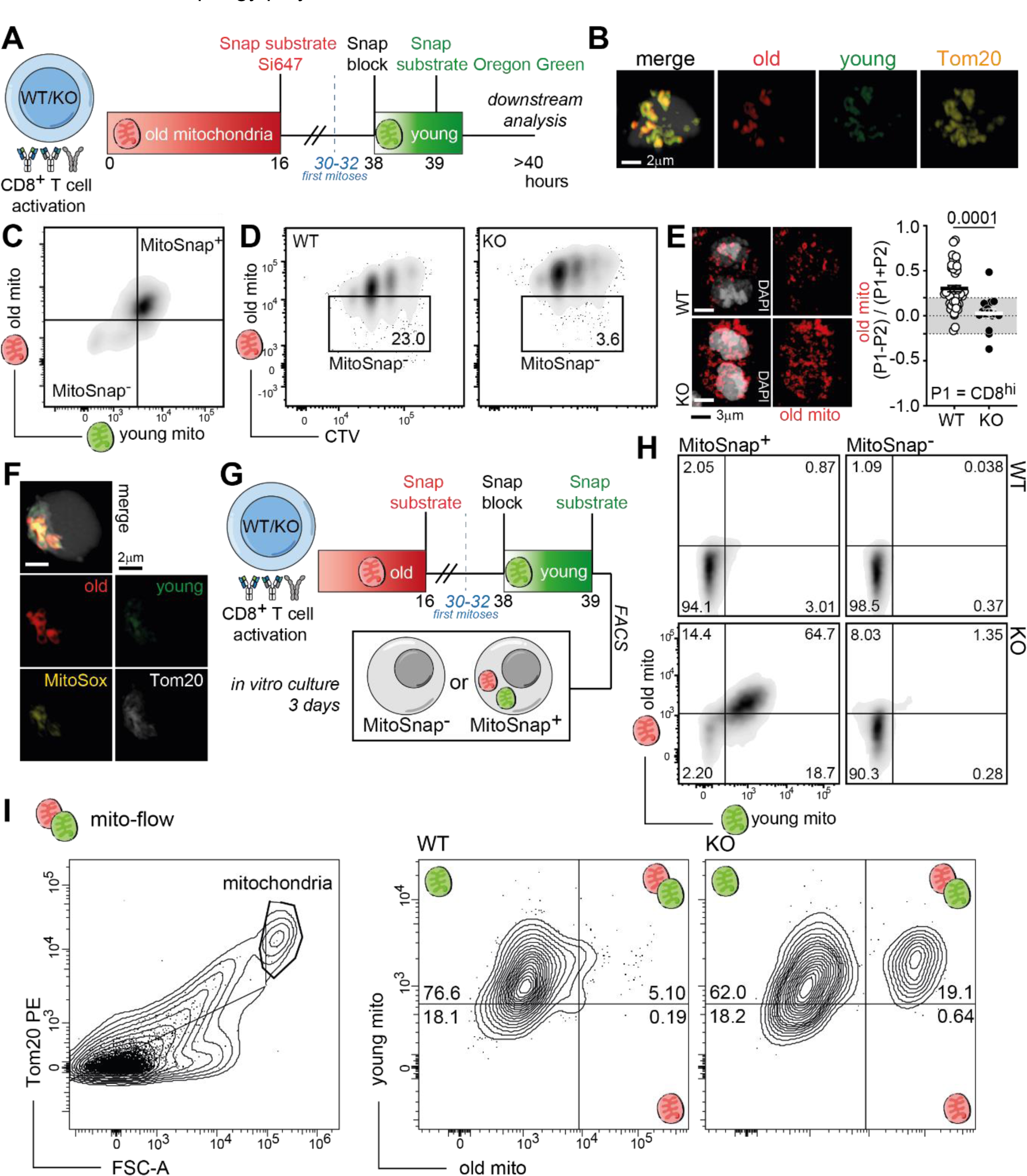
Inheritance of old mitochondria is autophagy-dependent. **(A)** Experimental layout: CTV labelled naïve MitoSnap CD8^+^ T cells (WT-*Atg16l1*^fl/-^ *Omp25*^fl/-^ *Ert2*^Cre^ or KO-*Atg16l1*^fl/fl^ *Omp25*^fl/-^ *Ert2*^Cre^) were activated on anti-CD3, anti-CD28 and Fc-ICAM-1 coated plates for 36-40 h. Cells were cultured in T cell medium containing 500 nM Z-4-Hydroxytamoxifen (4OHT). 16 h post-activation, cells were harvested and labelled with Snap-Cell 647-SiR to tag old mitochondria and cultured for a further 24 h, when Snap-Cell Block and Snap-Cell Oregon Green incubations allowed young organelle labelling. Downstream analysis was done >2 h after cell resting in complete T cell medium at 37°C. **(B)** Representative confocal microscopy images showing specificity of SnapTag labelling (staining overlaps with anti-Tom20 antibody labelling) in WT MitoSnap CD8^+^ T cells 36 h post-activation. **(C)** Representative flow cytometry plot of old and young mitochondria inheritance amongst activated MitoSnap CD8^+^ T cells following first cell division. **(D)** Representative flow cytometry plots showing inheritance of old mitochondria during several cell division cycles in both autophagy-sufficient (WT) and autophagy-deficient (KO) cells. **(E)** Representative confocal microscopy images of mitotic WT and KO MitoSnap CD8^+^ T cells 36-40 h post-activation. Asymmetric inheritance of old mitochondria was calculated in each group. Data are represented as mean ± SEM. Statistical analysis was performed using an unpaired two-tailed Student’s *t* test. Exact P values are depicted in the figure. **(F)** Representative confocal microscopy images showing overlap between MitoSOX staining and old mitochondria labelling in WT MitoSnap CD8^+^ T cells 36 h post-activation. **(G)** Experimental layout: CTV labelled MitoSnap CD8^+^ T cells (WT-*Atg16l1*^fl/-^ *Omp25*^fl/-^ *Ert2*^Cre^ or KO-*Atg16l1*^fl/fl^ *Omp25*^fl/-^ *Ert2*^Cre^) were activated and SnapTag labelled as in 2A. MitoSnap-cells and MitoSnap+ cells were sorted as depicted in 2C. Sorted cells were cultured for 3 days in T cell medium supplemented with IL-2, IL-7 and IL-15. **(H)** Representative plots from MitoSnap CD8^+^ T cells 3 days post sorting. Sorted MitoSnap-cells were used to set up gating strategy. **(I)** MitoSnap CD8^+^ T cells (WT-*Atg16l1*^fl/-^ *Omp25*^fl/-^ *Ert2*^Cre^ or KO-*Atg16l1*^fl/fl^ *Omp25*^fl/-^ *Ert2*^Cre^) were activated and SnapTag-labelled as in 2A. Mitochondria were purified and phenotyped by flow cytometry. Mitochondrial gating was determined based on size and Tom20 expression (left, also refer to Fig. S2B). SnapTag-labelling was preserved and maintenance of old and young organelle staining was evaluated in autophagy-sufficient and autophagy-deficient cells.

### Old mitochondria are cell fate determinants that impede memory CD8+ T cell differentiation

Next, we aimed to investigate whether the inheritance of aged mitochondria impacts the fate of CD8^+^ T cells *in vivo*. To achieve that, we generated OT-I CD45.1 MitoSnap mice. CD8^+^ T cells from OT-I mice express a transgenic TCR specific to OVA_257–264_ SIINFEKL peptide^37^. The transgenic TCR allows robust and specific TCR-activation of these cells and CD45.1 allows tracing of these cells in a host mouse. As antigen, we chose *Listeria monocytogenes* expressing OVA (LM-OVA) as an acute infection model. OT-I MitoSnap cells were activated *in vitro* and first-daughter cells sorted into MitoSnap+ and MitoSnap-populations. These two distinct populations of OT-I T cells were transferred to WT naïve CD45.2 C57BL/6 hosts and after >4 weeks host mice were infected with LM-OVA. The immune responses generated by the transferred OT-I MitoSnap cells were followed by blood kinetics and >4 weeks post-bacterial challenge (memory phase) we assessed the abundance, phenotype and function of remaining progenies (Fig. 3A). We observed that cell populations derived from originally MitoSnap-cells had superior ability to survive than those generated by from MitoSnap+ cells, as re-expansion potential upon LM-OVA infection was significantly higher in the first group (Fig. 3B). The higher frequencies of MitoSnap-progenies within the total CD8^+^ T cell population from the host were maintained throughout the course of the immune response. When spleens were analyzed at the memory phase, we confirmed that higher frequencies were also predictive of higher OT-I cell numbers (Fig. 3C). Upon in vitro re-stimulation, MitoSnap-progenies also produced more than twice as much IFNγ than their MitoSnap+ counterparts (Fig. 3D). We did not observe differences in the frequencies of KLRG1^-^CD127^+^ and KLRG1^+^CD127^-^ between MitoSnap+ and MitoSnap-progenies (Fig. S3A). *In vitro* Tat-Cre driven recombination of *Omp25*-SnapTag^fl/-^ CD8^+^ T cells resulted in similar results, i.e. MitoSnap-cells show higher re-expansion rates upon cognate antigen re-challenge than MitoSnap+ cells (Fig.S3B). Together, the phenotype and function of MitoSnap-CD8^+^ T cells suggest that they have better memory potential.

**Figure 3.**
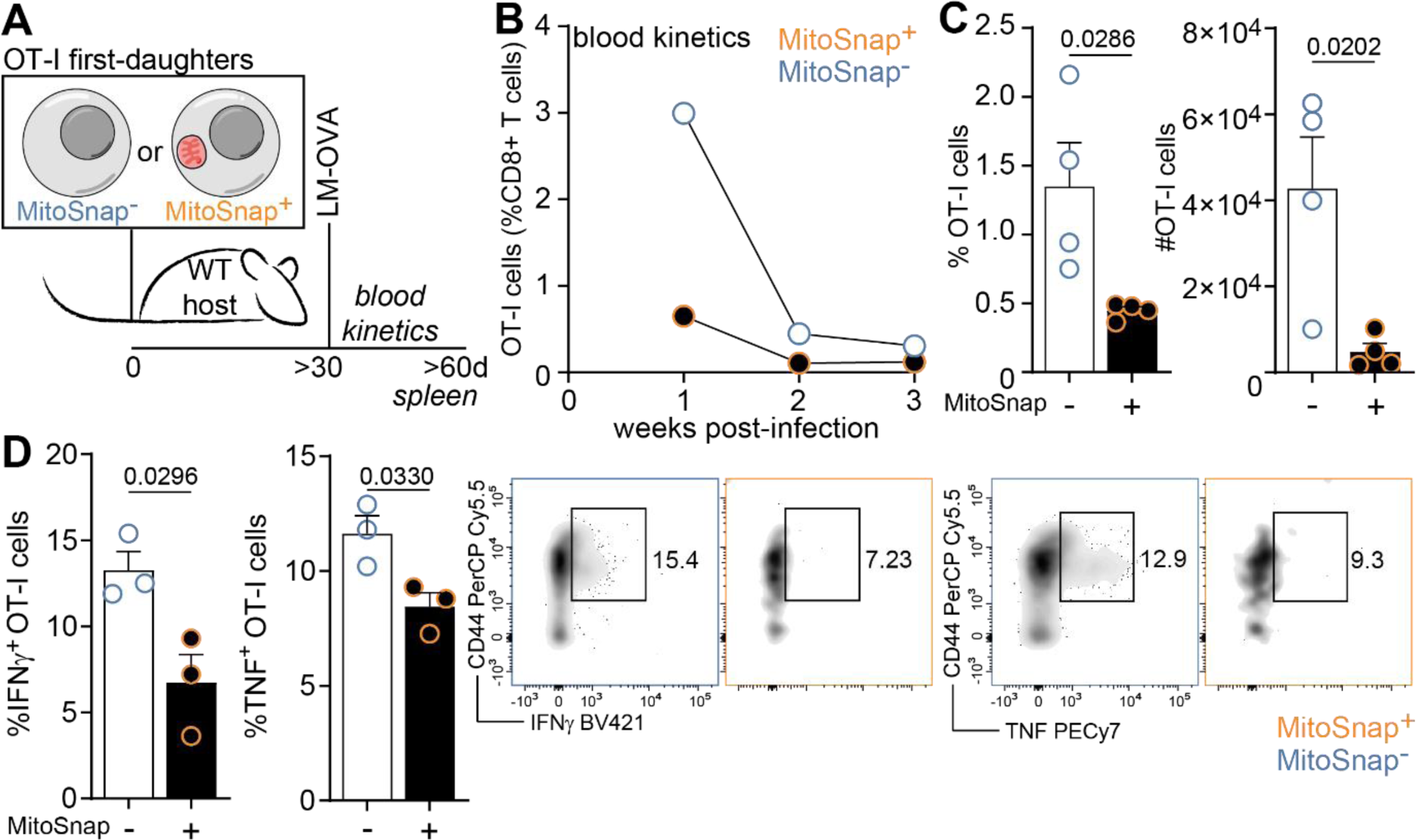
Old mitochondria are cell fate determinants that impede memory CD8^+^ T cell differentiation. **(A)** Experimental layout: CTV labelled naïve OT-I MitoSnap CD8^+^ T cells (*Atg16l1*^fl/-^ *Omp25*^fl/-^ *Ert2*^Cre^) were activated on anti-CD3, anti-CD28 and Fc-ICAM-1 coated plates for 36-40 h. Cells were cultured in T cell medium containing 500 nM Z-4-Hydroxytamoxifen (4OHT). 16 h post-activation, cells were harvested and labelled with Snap-Cell 647-SiR to tag old mitochondria and put back in culture. 24 h later cells were sorted into MitoSnap+ and MitoSnap-cells. 5×10^3^ cells were transferred to new hosts (CD45.1 and CD45.2 congenic markers were used to trace transferred cells). >30 days following adoptive cell transfer, host mice were infected with 2000 colony forming units (CFU) of *Listeria monocytogenes* expressing ovalbumin (OVA) (LM-OVA). Immune responses were evaluated in the blood and spleen. **(B)** Frequencies of OT-I cells within the CD8^+^ T-cell population in the blood. **(C)** Frequency and numbers of adoptively transferred OT-I cells within CD8^+^ T cells in the spleens of recipient mice. **(D)** Frequency of splenic IFNγ and TNF OT-I producing cells (left). Representative flow cytometry plots of MitoSnap+ and MitoSnap-cytokine producing cells (right). C, D: Data are represented as mean ± SEM. Statistical analysis was performed using an unpaired two-tailed Student’s *t* test. Exact P values are depicted in the figure. Representative data of 1 out of 4 experiments.

### Inheritance of aged mitochondria counteracts cellular quiescence

Effector CD8^+^ T cells are highly proliferative, while memory CD8^+^ T cells divide slower and are more quiescent. This is a feature that is established early on following T cell activation, with cell cycle speed predictive of CD8^+^ T cell fate^22,38^. CD8^+^ T cells with different clonal expansion rates have different metabolic demands, effector cells being more reliant on glycolysis, while long-lived naïve and memory cells mostly perform mitochondrial oxidative phosphorylation and fatty acid oxidation to produce ATP^32,33^. Thus, we investigated whether the metabolism of first-daughter CD8^+^ T cells is impacted by the inheritance of aged mitochondria using a modified version of the Scenith assay^39^. This assay allows measurement of metabolic dependencies by quantifying cellular translation rates, which highly correlate with ATP production. Translation is measured by the incorporation of O-propargyl-puromycin (OPP, a puromycin analogue), which can be visualized using click chemistry and flow cytometry. Metabolic reliance is evaluated by the addition of different inhibitors targeting glycolysis or OXPHOS. MitoSnap+ CD8^+^ T cells inheriting old mitochondria exhibited higher global translation rates and reliance on glycolysis than MitoSnap-cells, which were more metabolically quiescent (Fig. 4A). Because the resolution of this assay did not allow us to quantify mitochondrial function in MitoSnap-cells, we also directly measured oxygen consumption rates (OCR, Fig. 4B) and ATP synthesis (Fig. 4C) in both purified populations. Besides a trend of higher basal respiration in MitoSnap-cells, we did not observe significant differences between the two populations (Fig. S4A). We speculate that differences in mitochondrial respiration were not seen because defects in mitochondrial function might take longer than a timeline of 24 h, the time between old-organelle labelling and the experimental assay. We did the same metabolic measurements in autophagy-deficient MitoSnap+ and MitoSnap-cells, and obtained similar results (Fig 4B-C, S4A). To assess whether inheritance of distinct mitochondrial pools and differences in metabolic reliance cause differences in proliferation rates and survival, sorted MitoSnap+ and MitoSnap-cells were cultured in T cell medium containing IL-2, IL-7 and IL-15 in absence of T cell activation for further 3 or 7 days, respectively. CD8^+^ T cells inheriting aged mitochondria exhibited lower frequencies of slow-dividing cells, and more homogeneous proliferation profile in comparison to MitoSnap-cells, corroborating their less quiescent status that might contribute to precocious cell death (Fig. 4E). Autophagy-deficient cells showed slower proliferation rates independent of their mitochondrial inheritance profile, suggesting that autophagy loss might play a role in cell cycle arrest, which corroborates previous reports about the role of autophagy in degrading cyclin-dependent kinase inhibitor 1B (CDKN1B) in T cells^40^. Concerning survival in cytokine-limiting conditions, as expected for the effector population, MitoSnap+ cells showed lower viability than MitoSnap-cells after 7 days in culture (Fig. 4F). Interestingly, autophagy-sufficient remaining surviving cells exhibited distinct phenotypes, being CD44^+^CD62L^+^ cells, an expression pattern seen quiescent memory cells^22,38^, more abundant amongst MitoSnap-progenies (Fig. 4G). Autophagy-deficient cells did not exhibit differences in phenotype linked to early mitochondrial inheritance (Fig. S4B). Because in WT cells aged mitochondria are cleared after 3 days even in MitoSnap+ cells that inherited their mitochondria from the mother cell, our results suggest that old organelles inherited at first division counteract cellular quiescence at early stages post-T cell stimulation. In turn this promotes the emergence of a cell population that resembles short-lived effector CD8^+^ T cells. Our results provide the first unequivocal data linking organelle inheritance in mammals - here mitochondria - to changes in cell function that culminate in fate commitment of cells *in vivo*.

**Figure 4.**
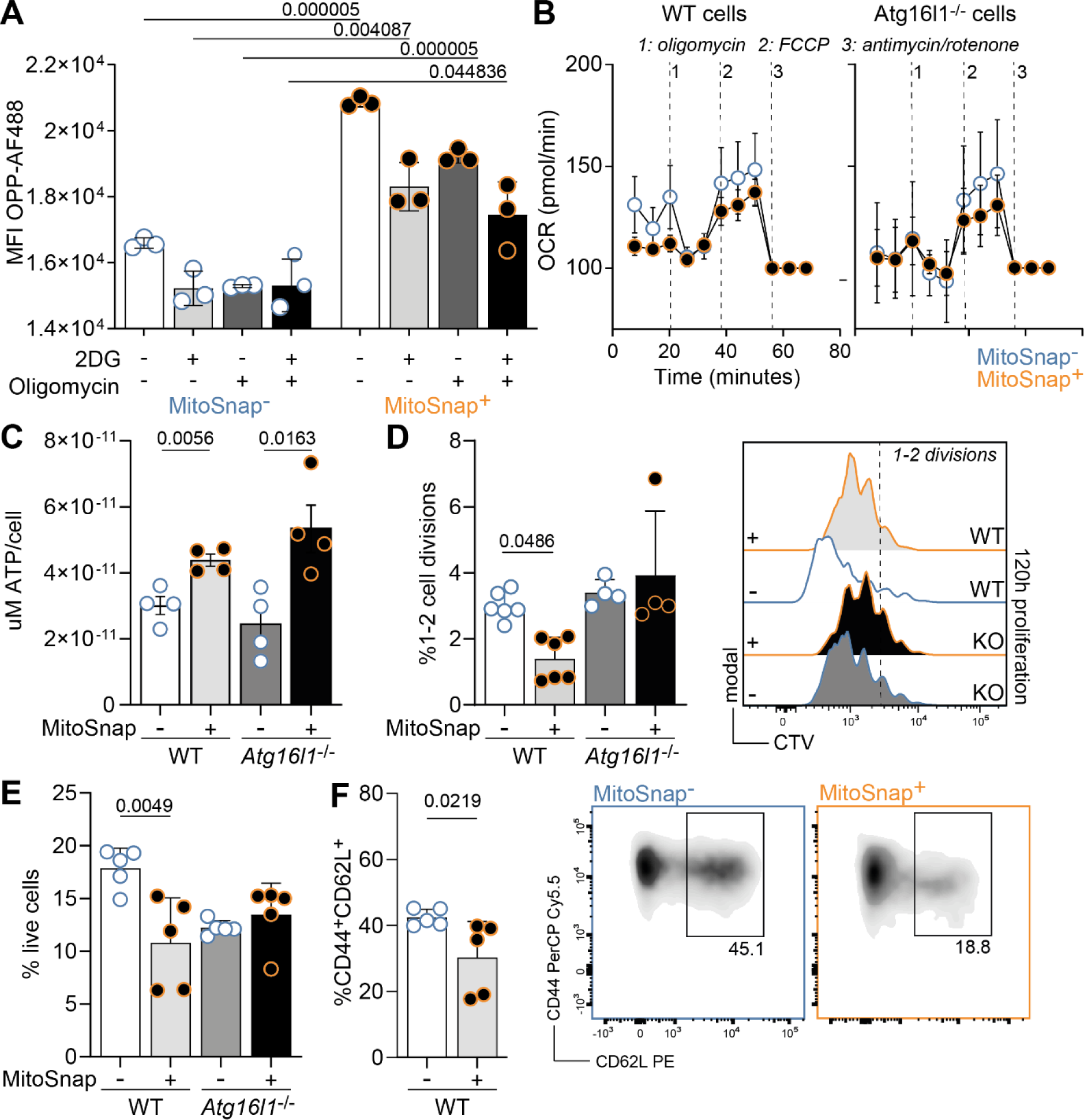
Inheritance of aged mitochondria counteracts cellular quiescence. **(A)** CTV labelled naïve MitoSnap CD8^+^ T cells (*Atg16l1*^fl/-^ *Omp25*^fl/-^ *Ert2*^Cre^) were activated on anti-CD3, anti-CD28 and Fc-ICAM-1 coated plates for 36-40 h. Cells were cultured in T cell medium containing 500 nM Z-4-Hydroxytamoxifen (4OHT). 16 h post-activation, cells were harvested and labelled with Snap-Cell 647-SiR (old mitochondria) and cultured for further 24 h, when cells were harvested and prepared for the Scenith assay to evaluate their metabolic reliance. OPP incorporation was used as a readout of translation. 2-Deoxy-D-glucose was used to inhibit glycolysis and oligomycin was used to inhibit mitochondrial respiration. A combination of both inhibitors was used to supress both metabolic pathways and obtain an OPP baseline. Analysis was done by flow cytometry, which allowed the discrimination of MitoSnap+ and MitoSnap-cells. Data are represented as mean ± SEM. Statistical analysis was performed using Two-Way ANOVA with Tukey’s post-hoc test. Exact P values are depicted in the figure. Representative data of 1 out of 3 experiments. **(B)** Oxygen consumption rate (OCR) of sorted MitoSnap+ and MitoSnap-first-daughter CD8^+^ T cells was measured under basal conditions and in response to indicated drugs. Data are represented as mean ± SEM. Datapoints represent 4 technical replicates from 2 biological samples. **(C)** ATP production by sorted MitoSnap+ and MitoSnap-first-daughter CD8^+^ T cells originally isolated from WT (*Atg16l1*^fl/-^ *Omp25*^fl/-^ *Ert2*^Cre^) or KO (*Atg16l1*^fl/fl^ *Omp25*^fl/-^ *Ert2*^Cre^) mice. Data are represented as mean ± SEM. Statistical analysis was performed using an unpaired two-tailed Student’s *t* test. Exact P values are depicted in the figure. Datapoints represent 4 technical replicates from 1 biological sample per group. Representative data from 1 out of 2 experiments. **(D)** WT and KO MitoSnap+ and MitoSnap-cells were sorted as represented in figure 2G and cultured for 3 days in T cell medium supplemented with IL-2, IL-7 and IL-15. Frequency of slow-dividing cells (1 or 2 divisions) was calculated. Data are represented as mean ± SEM. Statistical analysis was performed using One-Way ANOVA. Exact P values are depicted in the figure. Datapoints represent 1-3 technical replicates from 2 biological samples per group. Representative data from 1 out of 2 experiments. **(E)** WT and KO MitoSnap+ and MitoSnap-cells were sorted as represented in figure 2G and cultured for 7 days in T cell medium supplemented with IL-2, IL-7 and IL-15. Frequency of viable cells was calculated. Data are represented as mean ± SEM. Statistical analysis was performed using One-Way ANOVA. Exact P values are depicted in the figure. Datapoints represent 5 technical replicates from 1 biological sample per group. Representative data from 1 out of 2 experiments. **(F)** Frequency of CD44^+^ CD62L^+^ cells within surviving cells from E was calculated. Gating strategy is depicted (right panel). Data are represented as mean ± SEM. Statistical analysis was performed using an unpaired two-tailed Student’s *t* test. Exact P values are depicted in the figure. Datapoints represent 5 technical replicates from 1 biological sample per group. Representative data from 1 out of 2 experiments.

### Unequal inheritance of mitochondrial populations drives changes in the transcriptome and proteome of CD8+ T cells

Aiming to further identify the fate-divergency drivers found in cells and the metabolism of daughter cells inheriting distinct mitochondrial pools, we labelled old mitochondria in activated CD8^+^ T cells, sorted MitoSnap+ and MitoSnap-first-daughter cells and performed single-cell transcriptomics (scRNAseq) and bulk proteomics analysis of these two populations (Fig. 5A). Proteomics analysis of combined 4 experiments (6 samples per group) allowed us to identify a small list of differentially inherited proteins in these two populations. MitoSnap-cells expressed higher levels of Werner protein (WRN), an enzyme important for genome stability^41^, and NADH dehydrogenase 4 (mt-ND4), a protein involved in mitochondrial biogenesis as part of the mitochondrial respiratory chain complex I (gene ID: 4538, HGNC). MitoSnap+ cells were enriched in Hypoxia Inducible Factor 1 Subunit Alpha (HIF1a) and late endosomal/lysosomal adaptor 2 (LAMTOR2), proteins involved in mammalian target of rapamycin (mTOR) metabolism, which has been reported to boost effector CD8^+^ T cell differentiation^42,43^ (Fig. 5B). In other studies, including our own, transcriptional profiling of CD8^+^ T cell populations following one cycle of cell division was performed using bulk and single cell strategies^27–30,44^. However, these reports either relied on the expression of surface markers and reporter genes with the caveat of their dynamic expression to identify effector-like and memory-like cell daughters. They could not directly link the transcriptional divergences to asymmetric inheritance of cell fate determinants during mitosis, as cells were generated *in vivo* and could have emerged from both symmetric and asymmetric cell divisions. Unbiased clustering of single cell transcriptomes and visualization with uniform manifold approximation and projection (UMAP) plots, allowed us to define 15 clusters (Fig. 5C). Both types of cells were present in all clusters, but some were enriched in MitoSnap+ or MitoSnap-daughter-cells. By evaluating the expression of the genes encoding for proteomics enriched targets in our scRNAseq UMAP, we confirmed that there was a positive correlation between gene and protein expression. Furthermore, the MitoSnap+ proteome cluster was enriched in clusters 1 (lower-half), 2, 4 and 8-12, while the MitoSnap-proteome cluster was enriched in clusters 1 (upper-half), 3, 7 and 13 (Fig. 5D). Interestingly, these cluster regions matched MitoSnap+ and MitoSnap-enriched clusters concerning cell numbers (Fig. 5C). Based on the genes mostly expressed in each cluster, we could assign functional signatures to each of them. Clusters 1 and 5 exhibited very high expression of mitochondrial encoded genes. Clusters dominated by MitoSnap-cells showed a memory-related signature (Cluster 3), high expression of genes linked to mitochondrial function and biogenesis (Cluster 7) and redox balance (Cluster 13) or a transcriptional signature marked by genes involved in chemotaxis and adhesion (Cluster 6). Most of the other clusters had a majority of MitoSnap+ cells and exhibited transcriptional profiles that could be linked to effector functions (Clusters 8-12). These gene signatures are aligned with the functional readouts previously obtained, as MitoSnap-cells are the ones with higher mitochondrial turnover rates and memory potential, while MitoSnap+ cells are more proliferative and show lower survival rates in absence of TCR-stimulation, a feature of effector-like cells. We also selected genes extensively reported to promote effector or memory differentiation in CD8^+^ T cells and found that they were enriched in MitoSnap+ and MitoSnap-abundant clusters, respectively (Fig. S5A-C). Interestingly, we also found a cluster enriched in genes that are linked to one-carbon (1C) metabolism (e.g. *Mthfd2*, *Phgdh* and *Shmt2*, Cluster 2, Fig. S5D). This cluster is formed by a small majority of MitoSnap+ cells, but the 1C metabolism signature is stronger in this population in comparison to MitoSnap-cells (Fig. 5E). In CD4^+^ T cells 1C metabolism is essential for proliferation and effector function as an inducer of mTOR activity^45^. Thus, aiming to functionally validate this finding, we again measured metabolic reliance through Scenith using SHIN, an inhibitor of serine hydroxymethyltransferase (SHMT1/2) activity, a mitochondrial enzyme responsible for the catabolism of serine to glycine, key to one-carbon metabolism. Our results suggest that in MitoSnap+ cells SHIN1 treatment indeed suppresses their translation rates, a phenotype that was not shared by MitoSnap-cells (Fig. 5F). Taken together, these results further support that inheritance of mitochondrial pools of different ages determines T cell fate divergence and this is caused by distinct strategies to fulfil metabolic demands: MitoSnap-cells are more quiescent and quickly turn over mitochondria, which includes mitochondrial biogenesis, while MitoSnap+ cells keep old/damaged mitochondria, are more glycolytic and turn to one-carbon metabolism, one of the first consequences after a mitochondrial insult.

**Figure 5.**
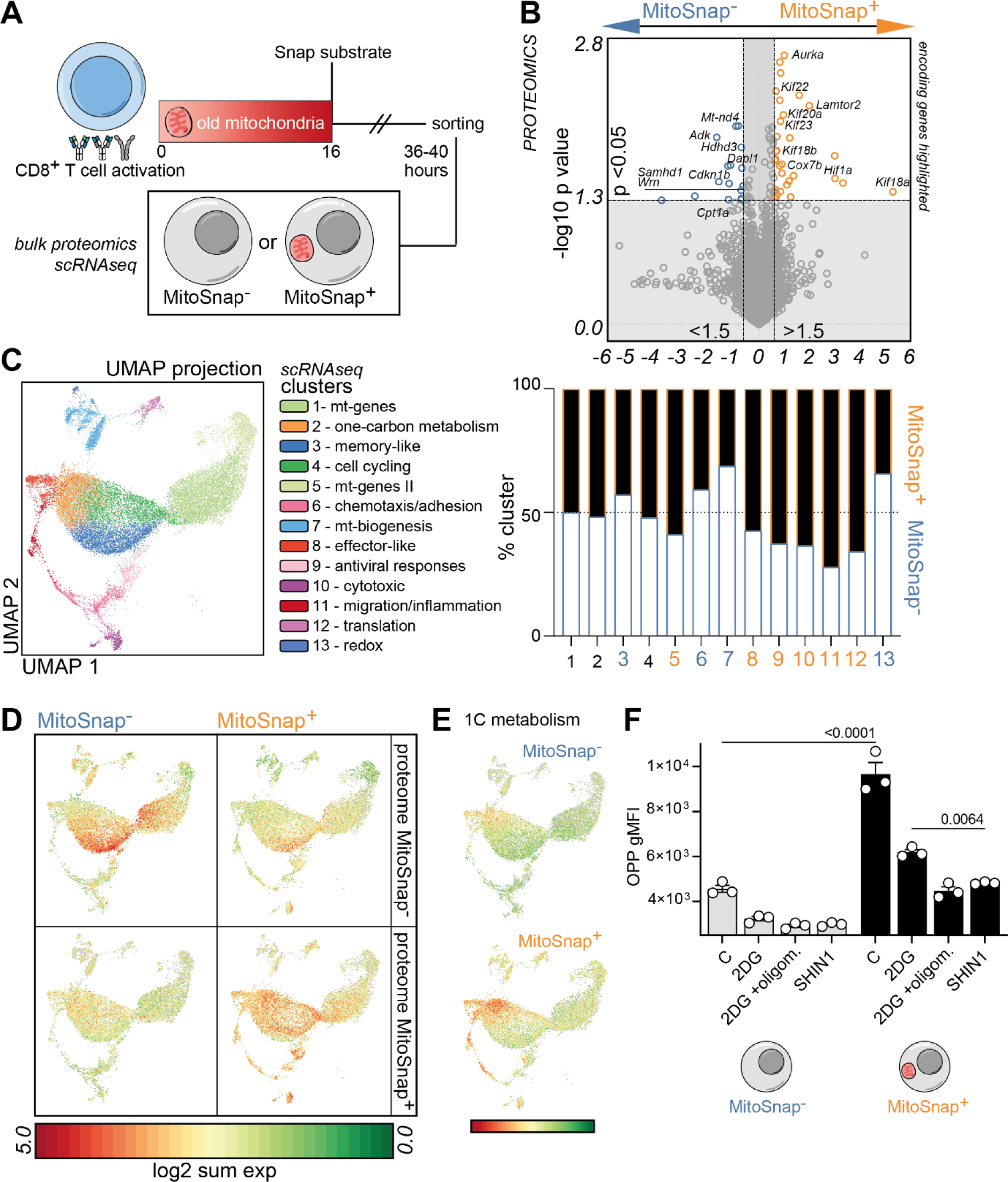
Unequal inheritance of mitochondrial populations drives changes in the transcriptome and proteome of CD8^+^ T cells. **(A)** Experimental layout: CTV labelled naïve MitoSnap CD8^+^ T cells (*Atg16l1*^fl/-^ *Omp25*^fl/-^ *Ert2*^Cre^) were activated on anti-CD3, anti-CD28 and Fc-ICAM-1 coated plates for 36-40 h. Cells were cultured in T cell medium containing 500 nM Z-4-Hydroxytamoxifen (4OHT). 16 h post-activation, cells were harvested and labelled with Snap-Cell 647-SiR (old mitochondria) and cultured for further 24 h. Cells were harvested and sorted into MitoSnap+ and MitoSnap-populations and their proteome and transcriptome were analyzed. **(B)** Volcano plot showing differentially inherited proteins by MitoSnap+ and MitoSnap-cells. Data pooled from 6 samples done in 4 independent experiments. MitoSnap+ and MitoSnap-enriched proteins (represented by their encoding genes) are highlighted in orange and blue, respectively. Proteomics volcano plot was done using Tableau. **(C)** UMAP and clustering of integrated MitoSnap+ and MitoSnap-cells obtained from 5 mice per group (left). Frequency of MitoSnap+ and MitoSnap-cells per cluster (right). **(D)** Genes encoding for proteins enriched in MitoSnap+ or MitoSnap- were projected onto UMAP clusters from 5C. **(E)** Genes involved in one-carbon (1C) metabolism were projected onto UMAP clusters. **(F)** CTV labelled naïve MitoSnap CD8^+^ T cells (*Atg16l1*^fl/-^ *Omp25*^fl/-^ *Ert2*^Cre^) were activated on anti-CD3, anti-CD28 and Fc-ICAM-1 coated plates for 36-40 h. Cells were cultured in T cell medium containing 500 nM Z-4-Hydroxytamoxifen (4OHT). 16 h post-activation, cells were harvested and labelled with Snap-Cell 647-SiR (old mitochondria) and cultured for further 24 h, when cells were harvested and prepared for the Scenith assay, aiming to evaluate their metabolic reliance. OPP incorporation was used as a readout of translation. 2-Deoxy-D-glucose was used to inhibit glycolysis and oligomycin was used to inhibit mitochondrial respiration. A combination of both inhibitors was used to supress both metabolic pathways and obtain an OPP baseline. SHIN1 was used to inhibit enzymes SHMT1/2. Analysis was done by flow cytometry, which allowed the discrimination of MitoSnap+ and MitoSnap-cells. Data are represented as mean ± SEM. Statistical analysis was performed using Two-Way ANOVA with Tukey’s post-hoc test. Exact P values are depicted in the figure. Representative data of 1 out of 2 experiments.

## Discussion

Most of the previous functional readouts evaluating the role of ACD in early fate decisions have relied on sorting daughter cells based on the expression of the surface marker CD8 or the transcription factor c-Myc, with CD8^hi^/c-Myc^hi^ cells being effector-like and CD8^lo^/c-Myc^lo^ cells being memory-like progenies^21,23,26^. However, the expression of these molecules is highly dynamic and does not necessarily result from asymmetric segregation events. A recent pioneering study used genetic barcoding to evaluate the transcriptome of genuine sister cells and demonstrated that early-fate trajectories can be established since first CD8^+^ T cell division^27^. However, overall there is no currently existing evidence to directly link this divergence to the inheritance of a fate determinant. Here we are first to show that asymmetric inheritance of pre-mitotic cell cargo causes divergent T cell fate commitment. This was possible because the MitoSnap system allows discrimination between events of inheritance and recent biogenesis. Tagging mitochondria before mitosis can be exclusively allocated to the pre-mitotic mother cell, thus guaranteeing that post-mitotic changes in cell phenotype do not interfere with its inheritance pattern, something which was not achieved in previous reports using expression of surface markers or reporter genes.

Mitochondria are organelles required to meet the cell’s energetic demands. They are the site of oxidative phosphorylation (OXPHOS), tricarboxylic acid (TCA) cycle and fatty acid oxidation (FAO), pathways involved in the generation of adenosine triphosphate (ATP). They are also involved in maintaining the redox balance of the cell, as they can produce reactive oxygen species (ROS), are involved in calcium signalling, can drive apoptotic cell death and, by being core metabolic modulators, also contribute to epigenetic regulation of cell function^5,46^. The results from several studies provide evidence that T cell fate is influenced by mitochondrial homeostasis, architecture and function: effector cells are highly glycolytic and memory cells rely on FAO^32,33,47,48^. Accordingly, mitochondrial quality control plays an important role in T cell fate decisions with mitophagy being a crucial regulator of cell survival^17,49,50^. Thus, mitochondria constitute a suitable cell cargo to be linked to differentiation trajectories, which was corroborated by our initial proteomics screening identifying mitochondrial-related proteins being differentially enriched in memory-like and effector-like CD8^+^ T cell daughters.

The emergence of cells that maintain or lose their MitoSnap labelling during CD8^+^ T cell proliferation could result from different cell biological processes and we dissected the mechanisms underlying the inheritance of mitochondria from the mother cell. Firstly, we identified that asymmetric cell division contributes to the polarized inheritance of old mitochondria. However, we observed that progenies able to clear old mitochondria also rapidly lost their labelling for young mitochondria, suggesting that MitoSnap-cells emerge from both segregation and degradation events, mitophagy levels being higher in this population. Autophagy and mitophagy support memory CD8^+^ T cell responses, but it remained unclear when these mechanisms are required to contribute to the formation of memory-precursors or the maintenance of long-lived cells^15–17^. To address whether autophagy plays a role in unequal mitochondrial inheritance, we used autophagy-deficient cells and found that asymmetric inheritance of old mitochondria was impaired and, as opposed to autophagy-competent cells, old mitochondria were kept for several days. These results and the symmetric proteome of CD8^hi^ and CD8^lo^ progenies from autophagy-deficient or old CD8^+^ T cells, corroborate our initial hypothesis and place ACD and autophagy/mitophagy as mechanisms that work synergistically to promote early asymmetric inheritance of cell fate determinants.

By following the frequencies of cells inheriting or not old mitochondria (MitoSnap+/-) over the course of the immune responses it became clear that MitoSnap-cells were more functional memory cells, as they showed better maintenance, re-expansion potential and ability to produce effector-cytokines upon re-stimulation. This resembles results obtained for CD8^hi^/c-Myc^hi^ and CD8^lo^/c-Myc^lo^ cells^21,26^, with the advantage that we can finally draw a definitive link between the inheritance of a cell cargo that already existed in the mother cell to the biased fate of its progenies. We then directed our attention to determine what drives the different fates of MitoSnap+ and MitoSnap-cells. By using sorted populations or approaches that provide single-cell resolution, we determined that the metabolism, survival and proliferative capacity of these progenies is different. Exhibiting lower translation rates, higher frequencies of slow-dividing cells and CD62L expression and better survival capacity in absence of antigen, MitoSnap-cells clearly showed a stronger memory phenotype than MitoSnap+ cells^22,38^. Although, surprisingly, we could not observe significant differences in mitochondrial respiration rates, MitoSnap+ cells relied more on glycolysis, a feature seen in effector CD8^+^ T cells. As old organelles also produced mitochondrial ROS as measured by MitoSOX, it is reasonable to assume that they have deteriorated mitochondrial fitness and that this might promote their early shift towards glycolysis^32^. Mitophagy has recently been reported to contribute to memory CD8^+^ T cell formation^17^. Our results add to this, showing that mitophagy contributes to the decision for memory CD8^+^ T cell fate commitment as early as the first mitosis following CD8^+^ T cell stimulation, as directly measured by the loss of young mitochondria generated after the first mitosis in MitoSnap-cells.

Finally, to obtain an unbiased overview of the differences between MitoSnap- and MitoSnap+ cells following the first mitosis post naïve CD8^+^ T cell activation, we performed both bulk proteomics and single cell transcriptomics of these two populations. In line with our expectations, we observed proteins linked to effector cell fate decision in MitoSnap+ cells and proteins linked to DNA health and mitochondrial biogenesis in MitoSnap-cells. It also came to our attention that a long list of kinesins (Kif genes) was enriched in effector-like MitoSnap+ daughters. Kinesins are motor proteins directly involved in intracellular trafficking of cell components along microtubules, which is important for organelle movement and for cell division events^51^, which fits with their less quiescent status and with the polarization of autophagosomes and mitochondria towards MitoSnap+ cells. Single cell transcriptomics allowed us to identify clusters that were enriched in MitoSnap+ and MitoSnap-cells. The presence of a memory-like cluster enriched in MitoSnap-cells, where this signature was stronger than in MitoSnap+ cells, further cements this cell type as the one inheriting the memory potential.

We became particularly interested in a cluster with a signature enriched in MitoSnap+ cells with higher expression of genes involved in 1C metabolism. 1C metabolism comprises methionine and folate cycles that provide 1C units to boost *de novo* synthesis of nucleotides and promote amino acid homeostasis and redox defence, particularly important in dividing cells such as cancer cells^52^. Enzymes involved in 1C metabolism can be found in the cytoplasm and the mitochondria, and both sets were upregulated in MitoSnap+ cells. Serine is an important donor of the 1C units when it is converted to glycine and in CD8^+^ T cells this amino acid has been shown to be important for clonal expansion of effector cells^53^. 1C metabolism has also been directly investigated in different CD4^+^ T cell subsets and results support its role in mTOR activation and the establishment of pro-inflammatory and highly proliferative populations^45^. Because expression of several amino acid transporters, including serine transporters, is upregulated in MitoSnap+ cells, which also exhibit defective translation upon C1 metabolism inhibition, our results provide further evidence of the role of this pathway as a regulator of cell fate decision.

Collectively, our results support the notion that organelle inheritance plays an important role in CD8^+^ T cell fate decision and contributes to the metabolic status of cell progenies. In cells from the haematopoietic lineage, the polarized presence of organelles during mitosis followed by long-term quantitative single-cell imaging has been reported, with the caveat that they were identified by dyes or probes that limit interpretation about their inheritance^54^. In CD8^+^ T cells, asymmetric mTOR activity in effector-like daughter cells has been linked to its translocation to lysosomes and amino acid sensing, but *in vivo* function readouts relied on correlative CD8 expression^43^. Concerning the asymmetric partitioning of degradation pathways, proteasome activity has been shown to contribute to distinct T-bet distribution between daughter cells, but results were not directly linked to *in vivo* T cell fates^24^. Here we show that organelle inheritance results from both degradation and segregation and that mitophagy and ACD work synergistically to form early memory-like cells and effector-like cells. As cells inheriting (or not) aged organelles are endowed with distinct metabolic signatures, our results suggest that therapeutic modulation of T cells can have different outcomes depending on when it is performed. Pre-mitotic modulation will globally impact on T cell differentiation, and post-mitotic approaches can selectively target a certain cell type, memory or effector, by inhibiting or improving its function. We anticipate that these findings will be relevant to a better understanding of how T cell diversity is early-imprinted and to foster the development of more efficient therapeutic strategies in the context of regenerative medicine and vaccination, which are particularly important in the context of ageing.

## Methods

### Study design

This study aimed to evaluate whether organelle inheritance controls CD8^+^ T cell differentiation. To achieve that, we investigated the role of asymmetric cell division and autophagy in patterns of mitochondria inheritance. The novel MitoSnap model was used to allow specific tracking of old vs. young organelles. We used imaging analysis of mitotic CD8^+^ T cells, flow cytometry readouts that allow single cell resolution, metabolic analysis and unbiased OMICS approaches to measure differences in phenotype and function between MitoSnap- and MitoSnap+ progenies. We used adoptive cell transfers of TCR-transgenic OT-I MitoSnap cells coupled to *Listeria monocytogenes*-OVA infections as a tool to assess immune responses and the impact of old organelle inheritance *in vivo*. All conclusions rely on at least two experiments. Every group consisted of at least two mice. No randomization or blinding was used.

### Animal models

All animal work was reviewed and approved by Oxford Ethical committee and the UK Home office under the project licenses PPL30/3388 and P01275425. Mice were bred under specific pathogen-free (SPF) conditions in-house, housed on a 12 h dark:light cycle, with a 30 min period of dawn and dusk and fed ad libitum. The temperature was kept between 20 and 24 °C, with a humidity level of 45–65%. Housing cages were individually ventilated and provided an enriched environment for the animals. MitoSnap mice (MGI:6466976; *Omp25*-SnapTag^fl/fl^) were kindly provided by the lab of Prof. Pekka Katajisto. This strain was then bred with CD45.1 *Atg16l1*^fl/fl^ *Ert2*^Cre^ OT-I mice expressing a TCR specific for OVA_257–264_ SIINFEKL peptide^37^, and maintained as CD45.1 or CD45.1/2 mice. Host mice in adoptive transfer experiments were either B6.SJL.CD45.1 or C57BL/6 naïve mice. Six-to-sixteen-week-old mice were considered young and > 100 week-old mice were considered aged.

### CD8^+^ T cell isolation and activation

Spleen and inguinal lymph nodes were harvested. Single-cell suspensions were used for naïve CD8^+^ T cell isolation using EasySep™ Mouse Naïve CD8^+^ T Cell Isolation Kit (Stemcell^TM^ Technologies) following manufacturer’s instructions. Purified populations were cultured (at 37°C, 5% CO_2_) in T cell medium: RPMI-1640 containing HEPES and l-glutamine (R5158, Sigma-Aldrich) supplemented with 10% filtered fetal bovine serum (Sigma-Aldrich), 1× Penicillin-Streptomycin (Sigma-Aldrich), 1× non-essential amino acids (Gibco), 50 μM β-mercaptoethanol (Gibco), and 1 mM sodium pyruvate (Gibco). T cell activation was done on anti-CD3 (5 μg/ml) (145-2C11, BioLegend), anti-CD28 (5 μg/ml) (37.51, BioLegend) and recombinant human or murine Fc-ICAM-1 (10 μg/ml) (R&D Systems) coated plates. 36-40h post activation cells were used in downstream assays. Autophagy deletion and/or SnapTag expression were induced by culturing cells in presence of 500 nM (Z)-4-hydroxytamoxifen (Sigma-Aldrich, H7904-5MG). To determine cell division events, cells were stained with CellTrace Violet^TM^ (Life Technologies) following manufacture’s guidelines.

### SnapTag labelling protocol

MitoSnap CD8^+^ T cells were labelled in 1 or 3 steps. Labelling of old organelles was done by harvesting CD8^+^ T cells 12-16 h post-activation and washing them in PBS (500 x*g*). Cells were incubated in T cell medium containing the first SnapSubstrate for 30 min at 37°C, washed in PBS and put back in culture in their original wells for further 20-24 h. When young organelle labelling was also performed, cells were harvested, washed and incubated with T cell medium containing 5 μM (Snap-Cell Block S9106S, New England Biolabs, NEB) for 30 min at 37°C. After washing, cells rested for 30-60 min in T cell medium and then incubated with the second SnapSubstrate for 30 min at 37°C. Fluorescent cell permeable Snap-Cell substrates (NEB) were used in the following concentrations: 3 μM (Snap-Cell 647-SiR S9102S), 3 μM (Snap-Cell TMR-Star S9105S), 5 μM (Snap-Cell Oregon Green S9104S).

### Cell survival and proliferation assays

Following activation, isolated MitoSnap CD8^+^ T cells (wild type vs. ATG16L1-deficient or MitoSnap+ vs. Mito Snap-first-daughter cells) were cultured in T cell medium supplemented with murine IL-2, IL-7 and IL-15 (5 ng/ml). Cell proliferation was evaluated 3 days later and cell survival was assessed 7 days later.

### Adoptive transfer and immunization

5-50×10^3^ FACS-purified MitoSnap+ or MitoSnap-cells (equal numbers in the same experiment to allow comparison between the two groups) were intravenously injected into naïve recipients. In the following day, mice were infected with 2×10^3^ colony-forming units (cfu) of *Listeria monocytogenes* expressing ovalbumin (LM-OVA) intravenously. LM-OVA was kindly provided by Prof. Audrey Gerard (Kennedy Institute of Rheumatology, University of Oxford). LM-OVA growth was done from frozen aliquots in Brain Heart Infusion (BHI) broth (Sigma, #53286-100G). Bacteria were used for infections when reaching exponential growth. Immune responses were tracked in the blood and at the memory phase spleens were harvested.

### Immunofluorescence staining and confocal microscopy

At different timepoints post-stimulation (pre-mitotic or mitotic/post-mitotic), CD8^+^ T cells were harvested. In some experiments, cells were incubated with 1-2 μM MitoSOX™ Mitochondrial Superoxide Indicator (Invitrogen) for 15 min at 37°C prior to harvesting. Cells were washed in PBS and transferred on Poly-L-Lysine (Sigma-Aldrich) treated coverslips, followed by incubation for 45-60 min at 37 °C. Attached cells were fixed with 2% methanol-free paraformaldehyde (PFA) in PBS (ThermoScientific) for 10 min, permeabilized with 0.3% Triton X-100 (Sigma-Aldrich) for 10 min and blocked in PBS containing 2% bovine serum albumin (BSA, Sigma-Aldrich) and 0.01% Tween 20 (Sigma-Aldrich) for 1 h at room temperature. The following antibodies were used to perform immunofluorescence stainings in murine cells: mouse anti-β-tubulin (Sigma-Aldrich), anti-mouse IgG AF488 (Abcam), anti-CD8 APC 53-6.7, BioLegend), anti-LC3B (D11) XP® Rabbit mAb PE (Cell Signalling). DAPI (Sigma-Aldrich) was used to detect DNA. ProLong™ Gold Antifade Mountant (ThermoScientific) was used as mounting medium. Mitotic cells (late anaphase to cytokinesis) were identified by nuclear morphology and/or presence of two microtubule organizing centres (MTOCs) and a clear tubulin bridge between two daughter cells. Forty to eighty Z-stacks (0.13μM) were acquired with a ZEISS 980 Airyscan 2 with a C-Apochromat 63x/1.2 W Corr magnification objective and the ZenBlue software. Data were analyzed using Fiji/ImageJ. Thresholds for quantification were setup individually for each fluorophore. Asymmetry rates were calculated based on the integrated density (volume and fluorescence intensity measurements were considered) of cell cargoes inherited by each daughter cell. This was done by using the following calculation: (P1-P2)/(P1+P2), where P1 is the daughter cell with higher integrated density of CD8 or old mitochondria. Any values above 0.2 or below −0.2 were considered asymmetric, which corresponds to one daughter-cell inheriting at least 1.5× more of a cell cargo than its sibling.

### Planar Supported Lipid Bilayers (PSLB)

Planar supported lipid bilayers were made as described previously ^55^. Briefly, glass coverslips were plasma-cleaned and assembled into disposable six-channel chambers (Ibidi). SLB were formed by incubation of each channel with small unilammellar vesicles containing 12.5 mol% 1,2-dioleoyl-sn-glycero-3-[(N-(5-amino-1-carboxypentyl) iminodiacetic acid) succinyl] (nickel salt) and 0.05 mol% 1,2-dioleoyl-sn-glycero-3-phosphoethanolamine-N-(cap biotinyl) (sodium salt) in 1,2-dioleoyl-sn-glycero-3-phosphocholine at total phospholipid concentration 0.4 mM. Chambers were filled with human serum albumin (HSA)-supplemented HEPES buffered saline (HBS), subsequently referred to as HBS/HSA. Following blocking with 5% casein in PBS containing 100 μM NiSO_4_, to saturate NTA sites, fluorescently labelled streptavidin was then coupled to biotin head groups. Biotinylated 2C11-fab fragments (30 molecules/μm^2^) and His-tagged ICAM-1 (200 molecules/μm^2^), and CD80 (100 molecules/μm^2^) were then incubated with the bilayers at concentrations to achieve the indicated site densities. Bilayers were continuous liquid disordered phase as determined by fluorescence recovery after photobleaching with a 10 µm bleach spot on an FV1200 confocal microscope (Olympus).

### T cell immunological synapse formation on PSLB

Naïve murine CD8^+^ T cells were incubated at 37°C on SLB. After 10 min, cells were fixed with 4% methanol-free formaldehyde in PHEM buffer (10 mM EGTA, 2 mM MgCl2, 60 mM Pipes, 25 mM HEPES, pH 7.0) and permeabilized with 0.1% Triton X-100 for 20 min at RT. Anti-CD3 staining was used to identify TCR regions and actin was labelled with fluorescent phalloidin.

### Total internal reflection fluorescence microscopy (TIRFM)

TIRFM was performed on an Olympus IX83 inverted microscope equipped with a 4-line (405 nm, 488 nm, 561 nm, and 640 nm laser) illumination system. The system was fitted with an Olympus UApON 150 × 1.45 numerical aperture objective, and a Photometrics Evolve Delta EMCCD camera to provide Nyquist sampling. Quantification of fluorescence intensity was performed with ImageJ.

### Flow Cytometry

Blood samples used for kinetics analysis were obtained from the tail vein at weeks 1, 2 and 3 post-LM-OVA challenge. At end-timepoints, spleens were harvested and single-cell splenocytes were prepared by meshing whole spleens through 70 µm strainers using a 1 ml syringe plunger. When cytokine production was assessed, splenocytes were incubated at 37 °C for 1 h with 1μM of SIINFEKL peptide, followed by 4 h in presence of SIINFEKL peptide + 10 µg/ml of brefeldin A (Sigma-Aldrich). Specific CD8^+^ T cells were evaluated by incubation with SIINFEKL_257-264_-APC-Labeled or SIINFEKL_257-264_-BV421-Labeled tetramers (NIH Tetramer Core Facility at Emory University). Erythrocytes were lysed by Red Blood Cell (RBC) Lysis buffer (Invitrogen). Conjugated antibodies used for surface staining were: anti-CD127 A7R34, anti-CD25 PC61 (AF700, PECy7, Biolegend; APC, eBioscience), anti-CD44 IM7 (AF700, BV785, PE, PerCPCy5.5, Biolegend), anti-CD45.1 A20 (BV785, FITC, PB, Biolegend), anti-CD45.2 104 (AF700, BV711, FITC, Biolegend), anti-CD62L MEL-14 (FITC, Biolegend; eF450, eBioscience), anti-KLRG1 2F1 (BV711, BV785, Biolegend), anti-CD8 53-6.7 (BV510, BV605, FITC, PE, Biolegend), anti-TCRβ H57-597 (APC-Cy7, PerCPCy5.5, Biolegend). Cells were incubated for 20 min at 4 °C. When intracellular staining was performed, cells were fixed/permed with 2× FACS Lysis Solution (BD Biosciences) with 0.08% Tween 20 (Sigma-Aldrich) for 10 min at RT, washed in PBS and incubated for 1h at RT with anti-IL2 JES6-5H4 (APC, Biolegend), anti-IFNγ XMG1.2 (BV421, Biolegend) and anti-TNF MP6-XT22 (PE-Cy7, ThermoFischer). Identification of viable cells was done by fixable near-IR dead cell staining (Life Technologies). All samples were washed and stored in PBS containing 2% FBS (Sigma-Aldrich) and 5 mM of EDTA (Sigma-Aldrich) before acquisition. Stained samples were acquired on a FACS LSR II (R/B/V) or a Fortessa X20 (R/B/V/YG) flow cytometer (BD Biosciences) with FACSDiva software. Data analysis was done using FlowJo software (FlowJo Enterprise, version 10.10, BD Biosciences).

### Cell sorting (FACS)

After activation, CTV- and SnapSubstrate-labelled MitoSnap CD8^+^ T cells were harvested and stained for phenotypical markers (anti-CD44 IM7, anti-CD45.1 A20, anti-CD45.2 104, anti-CD8 53-6.7 conjugated to different fluorophores depending on experiment, all Biolegend). Dead cells were excluded by staining cells with a fixable Live/Dead dye (Invitrogen, L34993 or L34957). Subpopulations of interest were sorted on a FACS Aria III cell sorter (BD Biosciences).

### Metabolic reliance measured by protein translation

We used a modified version of the Scenith assay^39^, which describes a high correlation between protein translation and ATP production. New protein synthesis was measured using the Click-iT Plus OPP Protein Synthesis Assay (Thermo Fisher, C10456) according to manufacturer’s protocol. In short, cells were incubated in T cell medium for 30 min at 37°C without any metabolic inhibitors or in presence of 1 μM oligomycin (Merck), 100 mM 2DG (Merck), a combination of both or 1 μM of SHIN1 (Cambridge Bioscience). This was followed by incubation with 10μM of alkynylated puromycin analog OPP for 30 min at 37°C. Click Chemistry was performed with Alexa Fluor 488™ dye picolyl azide. Metabolic reliance was assessed by comparing the OPP gMFI, used as an indicator of the relative translation rate, of inhibited samples to the vehicle control.

### Western Blot

Following (Z)-4-hydroxytamoxifen (Sigma-Aldrich, H7904-5MG) treatment for 24 h and/or bafilomycin A1 (BafA) treatment (10 nM) for 2 h or not, cells were washed with PBS and lysed in RIPA lysis buffer (Sigma-Aldrich) supplemented with complete Protease Inhibitor Cocktail (Roche) and PhosSTOP (Roche). Protein concentration was calculated by using the BCA Assay (ThermoFisher). Samples were diluted in 4x Laemmli Sample Buffer (Bio-Rad) and boiled at 100 °C for 5 min. 20 µg protein per sample were used for SDS-PAGE analysis. NuPAGE Novex 4%–12% Bis-Tris gradient gel (Invitrogen) with MOPS running buffer (Invitrogen) was used. Proteins were transferred to a PVDF membrane (Merck Millipore) and blocked with 5% skimmed milk-TBST (TBS 10x [Sigma-Aldrich] diluted to 1x in distilled water containing 0.1% Tween 20 [Sigma-Aldrich]) for 1h. Membranes were incubated at 4°C overnight with primary antibodies diluted in 1% skimmed milk-TBST and at room temperature for 1 h with secondary antibodies diluted in 1% skimmed milk-TBST supplemented 0.01% SDS. Primary antibodies used were: anti-ATG16L1, clone EPR15638 (Abcam, ab187671) and anti-GAPDH, clone 6C5 (Sigma-Aldrich, MAB374). Secondary antibodies used were: IRDye 680LT Goat anti-Mouse IgG (H + L) (Licor, 926-680-70) and IRDye 800CW Goat anti-Rabbit IgG (H + L) (Licor, 926-322-11). Images were acquired using the Odyssey CLx Imaging System. Data were analyzed using Image Studio Lite or Fiji.

### Mitochondrial isolation and flow cytometry (MitoFlow)

Autophagy-sufficient (*Atg16l1*^fl/-^ *Omp25*^fl/-^ *Ert2*^Cre^) and -deficient (*Atg16l1*^fl/fl^ *Omp25*^fl/-^ *Ert2*^Cre^) MitoSnap CD8^+^ T cells were activated, labelled for old (SNAP-Cell® TMR-Star, NEB) and young organelles (SNAP-Cell® Oregon Green, NEB), as previously described in the methods section, and after 40h washed with complete T cell medium. Cell pellets were resuspended in ice-cold mitochondria isolation buffer (320 mM sucrose, 2 mM EGTA,10 mM Tris-HCl, at pH 7.2 in water) and homogenized with a Dounce homogenizer with a 2 ml reservoir capacity (Abcam). We performed 20 strokes with a type B pretzel. The homogenizer was rinsed with distilled water before each sample was processed to avoid cross-contamination. Differential centrifugation of homogenates was done at 1,000 x*g* (4 °C for 8 min), which resulted in a pellet containing whole cells and isolated nuclei first. The supernatant containing the mitochondria was then transferred into new tubes and centrifuged at 17,000 x*g* (4 °C for 15 min). Enriched mitochondria, which appeared as brown-colored pellets, were fixed in 1% PFA in 0.5 ml PBS on ice for 15 min, followed by a wash with PBS. Mitochondria were resuspended in blocking buffer containing anti-Tom20-BV421 antibody for 20 min at RT. After washing with PBS, mitochondria were resuspended in 250 μl filtered (0.2 μm) PBS and acquired using a BD Fortessa X-20 flow cytometer. The threshold for SSC-A (log-scale) was set to the minimum value (20,000) to allow acquisition of subcellular particles. Submicron Particle Size Reference Beads (0.5 µm, 1 µm and 2 µm, Thermo Fisher Scientific) were also used to identify mitochondria.

### Metabolic flux analysis

MitoSnap+ and MitoSnap-cells were purified by FACS and their oxygen consumption rates (OCR) were measured using a XF96 MitoStress Test (Seahorse Agilent, 103015-100). Activated CD8^+^ T cells were washed in RPMI 1640 without sodium bicarbonate, 10 mM glucose, 1% FCS, 2 mM pyruvate and seeded in a XF plate (Agilent, 103793-100) coated with poly-L-lysine (Sigma-Aldrich) at equal densities in corresponding assay medium (XF Assay Medium, 103680-100) pH 7.4 supplemented with 10 mM glucose, 1 mM sodium pyruvate and 2 mM L-glutamine. Test compounds were sequentially injected to obtain the following concentrations: 1 µM oligomycin, 1.5 µM FCCP, 1 µM rotenone and 1 µM antimycin A. OCRs were normalized to cell number using CyQuant (Molecular Probes).

### ATP synthesis assay

Sorted MitoSnap+ and MitoSnap-CD8^+^ T cells were boiled in 100 mM Tris, 4 mM EDTA, pH 7.74 buffer for 2 min at 100°C. Following centrifugation, the supernatant was used for analysis. ATP levels were assessed using the ATP Bioluminescence Assay Kit CLS II (Roche) following the manufacturer’s instructions. The samples and ATP standard mixtures were swiftly combined with an equal volume of luciferase and promptly measured in a luminometer (BMG CLARIOstar Plus microplate reader). Normalization was performed by adjusting values based on the total number of sorted cells. Experiment was performed twice. Each experiment was done with 2 samples/group (each one pooled from 2 biological replicates) and at least four technical replicates per group.

### Proteomics

Proteomics analysis was done as previously described^56^. CD8^hi^ and CD8^lo^ or MitoSnap+ and MitoSnap-daughter-cells following naïve CD8^+^ T cell activation were purified by FACS. Cell pellets were washed 2× in PBS before being stored at −80°C prior to proteomics analysis. Samples were resuspended in 200 µl of S-Trap lysis buffer (10% SDS, 100mM Triethylammonium bicarbonate) and sonicated for 15 min (30 s on, 30 s off, 100% Amplitude, 70% Pulse). Samples were centrifuged and supernatants were transferred to fresh tubes. Protein quantification was done using the Micro BCA Protein Assay Kit (ThermoFisher). 150 μg of protein was processed using S-Trap mini columns (Protifi, #CO2-mini-80). The samples were digested overnight with 3.75 μg of trypsin (ThermoFisher, Pierce Trypsin Protease MS-Grade, #90057) with a second digest with the same amount of trypsin for 6 h the following day. Peptides were extracted, dried under vacuum and resuspended to 50 μl with 1% Formic Acid (ThermoFisher, #85178) and quantified using the Pierce Quantitative Fluorometric Peptide Assay (ThermoFisher, #23290).

Peptides were injected onto a nanoscale C18 reverse-phase chromatography system (UltiMate 3000 RSLC nano, ThermoFisher) and electrosprayed into an Orbitrap Exploris 480 Mass Spectrometer (MS) (ThermoFisher). For liquid chromatography the following buffers were used: buffer A (0.1% formic acid in Milli-Q water (v/v)) and buffer B (80% acetonitrile and 0.1% formic acid in Milli-Q water (v/v). Samples were loaded at 10 μL/min onto a trap column (100 μm × 2 cm, PepMap nanoViper C18 column, 5 μm, 100 Å, ThermoFisher) equilibrated in 0.1% trifluoroacetic acid (TFA). The trap column was washed for 3 min at the same flow rate with 0.1% TFA then switched in-line with a ThermoFisher, resolving C18 column (75 μm × 50 cm, PepMap RSLC C18 column, 2 μm, 100 Å). Peptides were eluted from the column at a constant flow rate of 300 nl/min with a linear gradient from 3% buffer B to 6% buffer B in 5 min, then from 6% buffer B to 35% buffer B in 115 min, and finally from 35% buffer B to 80% buffer B within 7 min. The column was then washed with 80% buffer B for 4 min. Two blanks were run between each sample to reduce carry-over. The column was kept at a constant temperature of 50°C. The data was acquired using an easy spray source operated in positive mode with spray voltage at 2.60 kV, and the ion transfer tube temperature at 250°C. The MS was operated in DIA mode. A scan cycle comprised a full MS scan (m/z range from 350-1650), with RF lens at 40%, AGC target set to custom, normalised AGC target at 300%, maximum injection time mode set to custom, maximum injection time at 20 ms, microscan set to 1 and source fragmentation disabled. MS survey scan was followed by MS/MS DIA scan events using the following parameters: multiplex ions set to false, collision energy mode set to stepped, collision energy type set to normalized, HCD collision energies set to 25.5, 27 and 30%, orbitrap resolution 30000, first mass 200, RF lens 40%, AGC target set to custom, normalized AGC target 3000%, microscan set to 1 and maximum injection time 55 ms. Data for both MS scan and MS/MS DIA scan events were acquired in profile mode.

Analysis of the DIA data was carried out using Spectronaut Biognosys, AG (version 14.7.201007.47784 for CD8^hi^ and CD8^lo^ cells obtained from young, *Atg16l1*-deficient and old mice; version 17.6.230428.55965 for MitoSnap+ and MitoSnap-cells). Data was analysed using the direct DIA workflow, with the following settings: imputation, profiling and cross run normalization were disabled; data Filtering to Qvalue; Precursor Qvalue Cutoff and Protein Qvalue Cutoff (Experimental) set to 0.01; maximum of 2 missed trypsin cleavages; PSM, Protein and Peptide FDR levels set to 0.01; cysteine carbamidomethylation set as fixed modification and acetyl (N-term), deamidation (asparagine, glutamine), oxidation of methionine set as variable modifications. The database used for CD8^hi^ and CD8^lo^ cells was mouse_swissprot_isoforms_extra_trembl_06_20.fasta (2020-06) and for mitosnap samples was the *Mus musculus* proteome obtained from uniprot.org (2022-02). Data filtering, protein copy number and concentration quantification was performed in the Perseus software package, version 1.6.6.0. Copy numbers were calculated using the proteomic ruler as described^31^. Samples were grouped according to the condition. P values were calculated via a two-tailed, unequal-variance t-test on log-normalized data. Elements with P values < 0.05 were considered significant, with a fold-change cut-off > 1.5 or < 0.67.

### Single cell transcriptomics

Single cell RNA sequencing libraries were prepared using the Chromium Single Cell 3’ GEX v3.1 assay (10X Genomics). In short, cell suspensions were encapsulated into Gel Beads in Emulsion (GEMs) using the Chromium Controller. Within each GEM, cell lysis and barcoded reverse transcription of RNA occurred, followed by cDNA amplification. The amplified cDNA underwent library construction via fragmentation, end-repair, A-tailing, adaptor ligation, and index PCR. Final libraries were sequenced on an Illumina NovaSeq 6000 system. Initial data processing was conducted with Cell Ranger 7.2.0.

Filtered output matrices were processed using Seurat. After loading the data and assigning unique identifiers to each dataset, cells with more than 30% mitochondrial gene content were excluded to ensure data quality (we used a less strict threshold because we were also interested in mitochondrial gene expression). The datasets were normalized using SCTransform, and PCA was conducted for dimensionality reduction. Integration of the datasets was achieved using the Harmony algorithm, followed by clustering and differential expression analysis. Finally, the integrated data were visualized using UMAP (down sampled to 13,000 cells per group). This methodology enabled a robust analysis while accounting for technical variations and maintaining biological integrity.

### Statistical analysis

To test if data point values were in a Gaussian distribution, a normality test was performed before applying parametric or non-parametric statistical analysis. When two groups were compared, unpaired Student’s t test or Mann-Whitney test were applied. When comparisons were done across more than two experimental groups, analysis were performed using One-Way ANOVA or Two-Way ANOVA with post hoc Tukey’s test multiple testing correction. P values were considered significant when < 0.05, and exact P values are provided in the figures. All analyses were done using GraphPad Prism 9 software.

## Data availability

The datasets generated or analyzed in this study are available from the corresponding lead author on reasonable request.

## Acknowledgments

We thank Dr. T. Youdale, Dr. L. Sinclair and Prof. D. Cantrell CBE, FRSE, FRS, FMedSci and the FingerPrints Proteomics Core Facility of the University of Dundee for their support with proteomics analysis. We thank T. Conrad, C. Fischer, C. Dietrich, F. Solinas and C. Braeuning from the BIH/MDC Genomics Platform for their support in generating the scRNAseq data. We thank E. Johnson (Dunn School, University of Oxford) for performing electron microscopy experiments. We thank P. C. Moreira, D. Andrew and M. Medghalchi (Kennedy Institute of Rheumatology BSU staff) for their support. We also thank L. Uhl for helping with LM-OVA infections. This work was funded by grants from the Wellcome Trust (Investigator award 103830/Z/14/Z and 220784/Z/20/Z to A.K.S., Sir Henry Wellcome Fellowship 220452/Z/20/Z to M.B., and PhD studentship award 203803/Z16/Z to F.C.R.), the Helmholz association (Helmholtz Distinguished Professorship Funding to recruit top-level international female scientists (W3) to A.K.S.), the European Union’s Horizon 2020 (Marie Sklodowska-Curie grant agreement number 893676 to M.B. and ERC-2021-SyG_951329 to E.C.B. and M.L.D.), the Swiss National Science Foundation (Early Postdoc.Mobility P2EZP3_188074 to M.B.), the European Molecular Biology Organization (EMBO LT postdoctoral Fellowship - ALTF1155-2019 to A.V.L.V.) and the Kennedy Trust for Rheumatology Research (KTTR) to Y.F.Y. and M.L.D. Flow cytometry and microscopy facilities were supported by KTTR.

## Authors contributions

M.B., A.V.L.V. and A.K.S., designed the experiments. M.B., A.V.L.V., E.B.C. and F.C.R. performed the experiments. H.B., M.L.D., P.K. provided expert assistance and guidance. M.B., A.V.L.V., A.H.K. analyzed the experiments. M.B. and A.K.S. wrote the manuscript.

## References

1. Stemberger, C., Huster, K.M., Koffler, M., Anderl, F., Schiemann, M., Wagner, H., and Busch, D.H. (2007). A single naive CD8+ T cell precursor can develop into diverse effector and memory subsets. Immunity 27, 985–997. 10.1016/j.immuni.2007.10.012.

2. Gerlach, C., Rohr, J.C., Perie, L., van Rooij, N., van Heijst, J.W., Velds, A., Urbanus, J., Naik, S.H., Jacobs, H., Beltman, J.B., et al. (2013). Heterogeneous differentiation patterns of individual CD8+ T cells. Science 340, 635–639. 10.1126/science.1235487.

3. Moller, S.H., Hsueh, P.C., Yu, Y.R., Zhang, L., and Ho, P.C. (2022). Metabolic programs tailor T cell immunity in viral infection, cancer, and aging. Cell Metab 34, 378–395. 10.1016/j.cmet.2022.02.003.

4. Chang, J.T., Wherry, E.J., and Goldrath, A.W. (2014). Molecular regulation of effector and memory T cell differentiation. Nat Immunol 15, 1104–1115. 10.1038/ni.3031.

5. Henning, A.N., Roychoudhuri, R., and Restifo, N.P. (2018). Epigenetic control of CD8(+) T cell differentiation. Nat Rev Immunol 18, 340–356. 10.1038/nri.2017.146.

6. Alsaleh, G., Panse, I., Swadling, L., Zhang, H., Richter, F.C., Meyer, A., Lord, J., Barnes, E., Klenerman, P., Green, C., and Simon, A.K. (2020). Autophagy in T cells from aged donors is maintained by spermidine and correlates with function and vaccine responses. Elife 9. 10.7554/eLife.57950.

7. Mittelbrunn, M., and Kroemer, G. (2021). Hallmarks of T cell aging. Nat Immunol 22, 687–698. 10.1038/s41590-021-00927-z.

8. Han, S., Georgiev, P., Ringel, A.E., Sharpe, A.H., and Haigis, M.C. (2023). Age-associated remodeling of T cell immunity and metabolism. Cell Metab 35, 36–55. 10.1016/j.cmet.2022.11.005.

9. Quinn, K.M., Fox, A., Harland, K.L., Russ, B.E., Li, J., Nguyen, T.H.O., Loh, L., Olshanksy, M., Naeem, H., Tsyganov, K., et al. (2018). Age-Related Decline in Primary CD8(+) T Cell Responses Is Associated with the Development of Senescence in Virtual Memory CD8(+) T Cells. Cell Rep 23, 3512–3524. 10.1016/j.celrep.2018.05.057.

10. Mogilenko, D.A., Shpynov, O., Andhey, P.S., Arthur, L., Swain, A., Esaulova, E., Brioschi, S., Shchukina, I., Kerndl, M., Bambouskova, M., et al. (2021). Comprehensive Profiling of an Aging Immune System Reveals Clonal GZMK(+) CD8(+) T Cells as Conserved Hallmark of Inflammaging. Immunity 54, 99–115 e112. 10.1016/j.immuni.2020.11.005.

11. Henson, S.M., Lanna, A., Riddell, N.E., Franzese, O., Macaulay, R., Griffiths, S.J., Puleston, D.J., Watson, A.S., Simon, A.K., Tooze, S.A., and Akbar, A.N. (2014). p38 signaling inhibits mTORC1-independent autophagy in senescent human CD8(+) T cells. J Clin Invest 124, 4004–4016. 10.1172/JCI75051.

12. Borsa, M., Barandun, N., Grabnitz, F., Barnstorf, I., Baumann, N.S., Pallmer, K., Baumann, S., Stark, D., Balaz, M., Oetiker, N., et al. (2021). Asymmetric cell division shapes naive and virtual memory T-cell immunity during ageing. Nat Commun 12, 2715. 10.1038/s41467-021-22954-y.

13. Sturmlechner, I., Jain, A., Mu, Y., Weyand, C.M., and Goronzy, J.J. (2023). T cell fate decisions during memory cell generation with aging. Semin Immunol 69, 101800. 10.1016/j.smim.2023.101800.

14. Clarke, A.J., and Simon, A.K. (2019). Autophagy in the renewal, differentiation and homeostasis of immune cells. Nat Rev Immunol 19, 170–183. 10.1038/s41577-018-0095-2.

15. Puleston, D.J., Zhang, H., Powell, T.J., Lipina, E., Sims, S., Panse, I., Watson, A.S., Cerundolo, V., Townsend, A.R., Klenerman, P., and Simon, A.K. (2014). Autophagy is a critical regulator of memory CD8(+) T cell formation. Elife 3. 10.7554/eLife.03706.

16. Xu, X., Araki, K., Li, S., Han, J.H., Ye, L., Tan, W.G., Konieczny, B.T., Bruinsma, M.W., Martinez, J., Pearce, E.L., et al. (2014). Autophagy is essential for effector CD8(+) T cell survival and memory formation. Nat Immunol 15, 1152–1161. 10.1038/ni.3025.

17. Franco, F., Bevilacqua, A., Wu, R.M., Kao, K.C., Lin, C.P., Rousseau, L., Peng, F.T., Chuang, Y.M., Peng, J.J., Park, J., et al. (2023). Regulatory circuits of mitophagy restrict distinct modes of cell death during memory CD8(+) T cell formation. Sci Immunol 8, eadf7579. 10.1126/sciimmunol.adf7579.

18. Pua, H.H., Guo, J., Komatsu, M., and He, Y.W. (2009). Autophagy is essential for mitochondrial clearance in mature T lymphocytes. J Immunol 182, 4046–4055. 10.4049/jimmunol.0801143.

19. Sunchu, B., and Cabernard, C. (2020). Principles and mechanisms of asymmetric cell division. Development 147. 10.1242/dev.167650.

20. Cobbold, S.P., Adams, E., Howie, D., and Waldmann, H. (2018). CD4(+) T Cell Fate Decisions Are Stochastic, Precede Cell Division, Depend on GITR Co-Stimulation, and Are Associated With Uropodium Development. Front Immunol 9, 1381. 10.3389/fimmu.2018.01381.

21. Chang, J.T., Palanivel, V.R., Kinjyo, I., Schambach, F., Intlekofer, A.M., Banerjee, A., Longworth, S.A., Vinup, K.E., Mrass, P., Oliaro, J., et al. (2007). Asymmetric T lymphocyte division in the initiation of adaptive immune responses. Science 315, 1687–1691. 10.1126/science.1139393.

22. Grabnitz, F., Stark, D., Shlesinger, D., Petkidis, A., Borsa, M., Yermanos, A., Carr, A., Barandun, N., Wehling, A., Balaz, M., et al. (2023). Asymmetric cell division safeguards memory CD8 T cell development. Cell Rep 42, 112468. 10.1016/j.celrep.2023.112468.

23. King, C.G., Koehli, S., Hausmann, B., Schmaler, M., Zehn, D., and Palmer, E. (2012). T cell affinity regulates asymmetric division, effector cell differentiation, and tissue pathology. Immunity 37, 709–720. 10.1016/j.immuni.2012.06.021.

24. Chang, J.T., Ciocca, M.L., Kinjyo, I., Palanivel, V.R., McClurkin, C.E., Dejong, C.S., Mooney, E.C., Kim, J.S., Steinel, N.C., Oliaro, J., et al. (2011). Asymmetric proteasome segregation as a mechanism for unequal partitioning of the transcription factor T-bet during T lymphocyte division. Immunity 34, 492–504. 10.1016/j.immuni.2011.03.017.

25. Pollizzi, K.N., Sun, I.H., Patel, C.H., Lo, Y.C., Oh, M.H., Waickman, A.T., Tam, A.J., Blosser, R.L., Wen, J., Delgoffe, G.M., and Powell, J.D. (2016). Asymmetric inheritance of mTORC1 kinase activity during division dictates CD8(+) T cell differentiation. Nat Immunol 17, 704–711. 10.1038/ni.3438.

26. Verbist, K.C., Guy, C.S., Milasta, S., Liedmann, S., Kaminski, M.M., Wang, R., and Green, D.R. (2016). Metabolic maintenance of cell asymmetry following division in activated T lymphocytes. Nature 532, 389–393. 10.1038/nature17442.

27. Liedmann, S., Liu, X., Guy, C.S., Crawford, J.C., Rodriguez, D.A., Kuzuoglu-Ozturk, D., Guo, A., Verbist, K.C., Temirov, J., Chen, M.J., et al. (2022). Localization of a TORC1-eIF4F translation complex during CD8(+) T cell activation drives divergent cell fate. Mol Cell 82, 2401–2414 e2409. 10.1016/j.molcel.2022.04.016.

28. Borsa, M., Barnstorf, I., Baumann, N.S., Pallmer, K., Yermanos, A., Grabnitz, F., Barandun, N., Hausmann, A., Sandu, I., Barral, Y., and Oxenius, A. (2019). Modulation of asymmetric cell division as a mechanism to boost CD8(+) T cell memory. Sci Immunol 4. 10.1126/sciimmunol.aav1730.

29. Metz, P.J., Arsenio, J., Kakaradov, B., Kim, S.H., Remedios, K.A., Oakley, K., Akimoto, K., Ohno, S., Yeo, G.W., and Chang, J.T. (2015). Regulation of asymmetric division and CD8+ T lymphocyte fate specification by protein kinase Czeta and protein kinase Clambda/iota. J Immunol 194, 2249–2259. 10.4049/jimmunol.1401652.

30. Quezada, L.K., Jin, W., Liu, Y.C., Kim, E.S., He, Z., Indralingam, C.S., Tysl, T., Labarta-Bajo, L., Wehrens, E.J., Jo, Y., et al. (2023). Early transcriptional and epigenetic divergence of CD8+ T cells responding to acute versus chronic infection. PLoS Biol 21, e3001983. 10.1371/journal.pbio.3001983.

31. Wisniewski, J.R., Hein, M.Y., Cox, J., and Mann, M. (2014). A "proteomic ruler" for protein copy number and concentration estimation without spike-in standards. Mol Cell Proteomics 13, 3497–3506. 10.1074/mcp.M113.037309.

32. Buck, M.D., O’Sullivan, D., Klein Geltink, R.I., Curtis, J.D., Chang, C.H., Sanin, D.E., Qiu, J., Kretz, O., Braas, D., van der Windt, G.J., et al. (2016). Mitochondrial Dynamics Controls T Cell Fate through Metabolic Programming. Cell 166, 63–76. 10.1016/j.cell.2016.05.035.

33. Pearce, E.L., Walsh, M.C., Cejas, P.J., Harms, G.M., Shen, H., Wang, L.S., Jones, R.G., and Choi, Y. (2009). Enhancing CD8 T-cell memory by modulating fatty acid metabolism. Nature 460, 103–107. 10.1038/nature08097.

34. Emurla, H., Barral, Y., and Oxenius, A. (2021). Role of mitotic diffusion barriers in regulating the asymmetric division of activated CD8 T cells. bioRxiv, 2021.2009.2010.458880. 10.1101/2021.09.10.458880.

35. Keppler, A., Gendreizig, S., Gronemeyer, T., Pick, H., Vogel, H., and Johnsson, K. (2003). A general method for the covalent labeling of fusion proteins with small molecules in vivo. Nat Biotechnol 21, 86–89. 10.1038/nbt765.

36. Katajisto, P., Dohla, J., Chaffer, C.L., Pentinmikko, N., Marjanovic, N., Iqbal, S., Zoncu, R., Chen, W., Weinberg, R.A., and Sabatini, D.M. (2015). Stem cells. Asymmetric apportioning of aged mitochondria between daughter cells is required for stemness. Science 348, 340–343. 10.1126/science.1260384.

37. Hogquist, K.A., Jameson, S.C., Heath, W.R., Howard, J.L., Bevan, M.J., and Carbone, F.R. (1994). T cell receptor antagonist peptides induce positive selection. Cell 76, 17–27. 10.1016/0092-8674(94)90169-4.

38. Plambeck, M., Kazeroonian, A., Loeffler, D., Kretschmer, L., Salinno, C., Schroeder, T., Busch, D.H., Flossdorf, M., and Buchholz, V.R. (2022). Heritable changes in division speed accompany the diversification of single T cell fate. Proc Natl Acad Sci U S A 119. 10.1073/pnas.2116260119.

39. Arguello, R.J., Combes, A.J., Char, R., Gigan, J.P., Baaziz, A.I., Bousiquot, E., Camosseto, V., Samad, B., Tsui, J., Yan, P., et al. (2020). SCENITH: A Flow Cytometry-Based Method to Functionally Profile Energy Metabolism with Single-Cell Resolution. Cell Metab 32, 1063–1075 e1067. 10.1016/j.cmet.2020.11.007.

40. Jia, W., He, M.X., McLeod, I.X., Guo, J., Ji, D., and He, Y.W. (2015). Autophagy regulates T lymphocyte proliferation through selective degradation of the cell-cycle inhibitor CDKN1B/p27Kip1. Autophagy 11, 2335–2345. 10.1080/15548627.2015.1110666.

41. Pichierri, P., Ammazzalorso, F., Bignami, M., and Franchitto, A. (2011). The Werner syndrome protein: linking the replication checkpoint response to genome stability. Aging (Albany NY) 3, 311–318. 10.18632/aging.100293.

42. Araki, K., Turner, A.P., Shaffer, V.O., Gangappa, S., Keller, S.A., Bachmann, M.F., Larsen, C.P., and Ahmed, R. (2009). mTOR regulates memory CD8 T-cell differentiation. Nature 460, 108–112. 10.1038/nature08155.

43. Pollizzi, K.N., Patel, C.H., Sun, I.H., Oh, M.H., Waickman, A.T., Wen, J., Delgoffe, G.M., and Powell, J.D. (2015). mTORC1 and mTORC2 selectively regulate CD8(+) T cell differentiation. J Clin Invest 125, 2090–2108. 10.1172/JCI77746.

44. Kakaradov, B., Arsenio, J., Widjaja, C.E., He, Z., Aigner, S., Metz, P.J., Yu, B., Wehrens, E.J., Lopez, J., Kim, S.H., et al. (2017). Early transcriptional and epigenetic regulation of CD8(+) T cell differentiation revealed by single-cell RNA sequencing. Nat Immunol 18, 422–432. 10.1038/ni.3688.

45. Sugiura, A., Andrejeva, G., Voss, K., Heintzman, D.R., Xu, X., Madden, M.Z., Ye, X., Beier, K.L., Chowdhury, N.U., Wolf, M.M., et al. (2022). MTHFD2 is a metabolic checkpoint controlling effector and regulatory T cell fate and function. Immunity 55, 65–81 e69. 10.1016/j.immuni.2021.10.011.

46. Franco, F., Jaccard, A., Romero, P., Yu, Y.R., and Ho, P.C. (2020). Metabolic and epigenetic regulation of T-cell exhaustion. Nat Metab 2, 1001–1012. 10.1038/s42255-020-00280-9.

47. van der Windt, G.J., Everts, B., Chang, C.H., Curtis, J.D., Freitas, T.C., Amiel, E., Pearce, E.J., and Pearce, E.L. (2012). Mitochondrial respiratory capacity is a critical regulator of CD8+ T cell memory development. Immunity 36, 68–78. 10.1016/j.immuni.2011.12.007.

48. O’Sullivan, D., van der Windt, G.J., Huang, S.C., Curtis, J.D., Chang, C.H., Buck, M.D., Qiu, J., Smith, A.M., Lam, W.Y., DiPlato, L.M., et al. (2014). Memory CD8(+) T cells use cell-intrinsic lipolysis to support the metabolic programming necessary for development. Immunity 41, 75–88. 10.1016/j.immuni.2014.06.005.

49. Corrado, M., Samardzic, D., Giacomello, M., Rana, N., Pearce, E.L., and Scorrano, L. (2021). Deletion of the mitochondria-shaping protein Opa1 during early thymocyte maturation impacts mature memory T cell metabolism. Cell Death Differ 28, 2194–2206. 10.1038/s41418-021-00747-6.

50. Girotra, M., Chiang, Y.H., Charmoy, M., Ginefra, P., Hope, H.C., Bataclan, C., Yu, Y.R., Schyrr, F., Franco, F., Geiger, H., et al. (2023). Induction of mitochondrial recycling reverts age- associated decline of the hematopoietic and immune systems. Nat Aging 3, 1057–1066. 10.1038/s43587-023-00473-3.

51. Endow, S.A., Kull, F.J., and Liu, H. (2010). Kinesins at a glance. J Cell Sci 123, 3420–3424. 10.1242/jcs.064113.

52. Ducker, G.S., and Rabinowitz, J.D. (2017). One-Carbon Metabolism in Health and Disease. Cell Metab 25, 27–42. 10.1016/j.cmet.2016.08.009.

53. Ma, E.H., Bantug, G., Griss, T., Condotta, S., Johnson, R.M., Samborska, B., Mainolfi, N., Suri, V., Guak, H., Balmer, M.L., et al. (2017). Serine Is an Essential Metabolite for Effector T Cell Expansion. Cell Metab 25, 345–357. 10.1016/j.cmet.2016.12.011.

54. Loeffler, D., Schneiter, F., Wang, W., Wehling, A., Kull, T., Lengerke, C., Manz, M.G., and Schroeder, T. (2022). Asymmetric organelle inheritance predicts human blood stem cell fate. Blood 139, 2011–2023. 10.1182/blood.2020009778.

55. Alpert, A., Pickman, Y., Leipold, M., Rosenberg-Hasson, Y., Ji, X., Gaujoux, R., Rabani, H., Starosvetsky, E., Kveler, K., Schaffert, S., et al. (2019). A clinically meaningful metric of immune age derived from high-dimensional longitudinal monitoring. Nat Med 25, 487–495. 10.1038/s41591-019-0381-y.

56. Jenkins, B.J., Blagih, J., Ponce-Garcia, F.M., Canavan, M., Gudgeon, N., Eastham, S., Hill, D., Hanlon, M.M., Ma, E.H., Bishop, E.L., et al. (2023). Canagliflozin impairs T cell effector function via metabolic suppression in autoimmunity. Cell Metab 35, 1132–1146 e1139. 10.1016/j.cmet.2023.05.001.

