## Supplementary Figures for "Inheritance of old mitochondria controls early CD8^+^ T cell fate commitment and is regulated by autophagy"

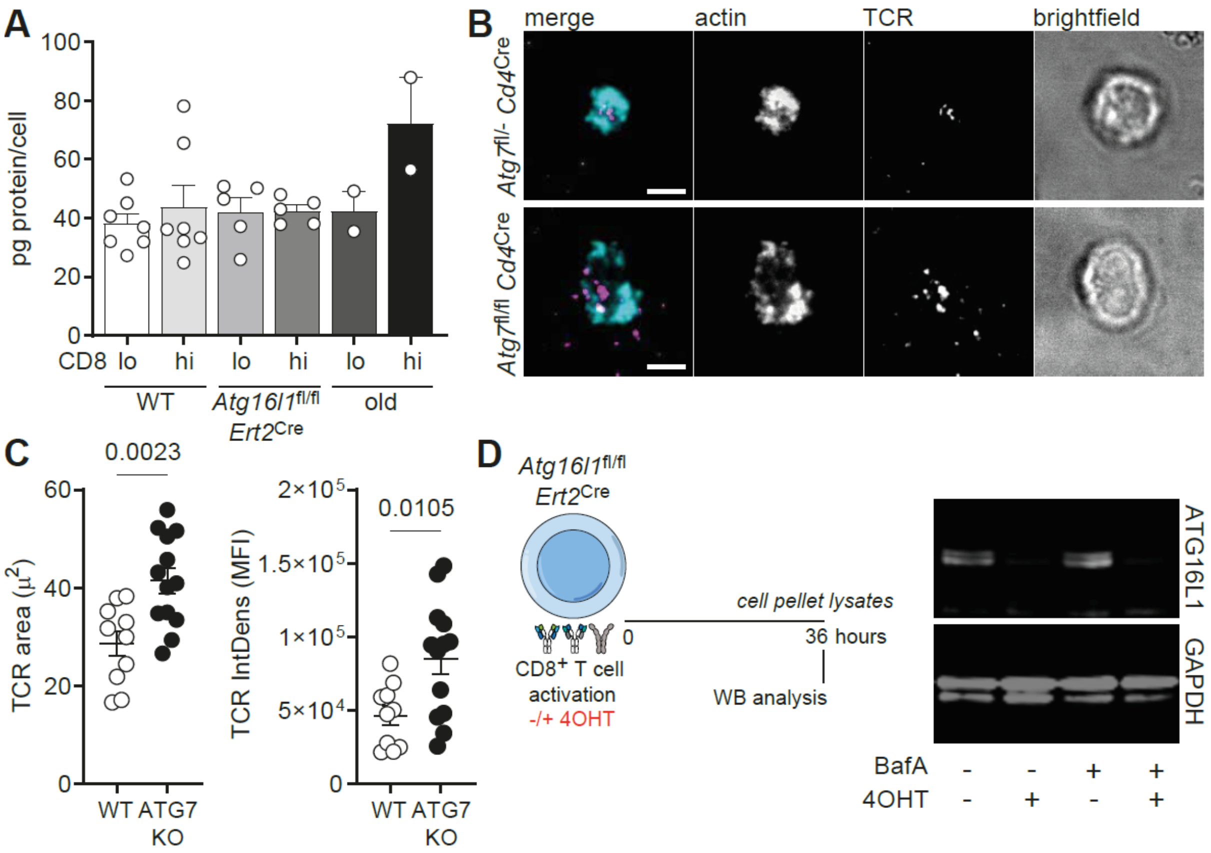


**Figure S1. Constitutive autophagy-deficiency changes immune synapse architecture, which makes inducible-deletion of *Atg16l1* a more suitable model to evaluate the impact of asymmetric inheritance in CD8⁺ T cells. (A)** $CD8^{hi}$ and $CD8^{lo}$ first-daughter cells were sorter and sent for proteomics analysis. Protein concentration per cell was comparable across groups. Data are represented as mean ± SEM. **(B)** Representative TIRF-images of immunological synapses of autophagy-sufficient and -deficient cells. Naïve $CD8^{+}$ T cells were added on a planar supported lipid bilayer (PSLB) containing anti-TCR, ICAM-1, and CD80. Cells were fixed after 10 min and stained for CD3 and actin. Scale bar represents 1μm. **(C)** Measurements of TCR area and intensity at immunological synapses. Data are represented as mean ± SEM. Statistical analysis was performed using an unpaired two-tailed Student's *t* test. Exact P values are depicted in the figure. **(D)** *Atg16l1*$^{fl/fl}$ *Ert2*$^{Cre}$ $CD8^{+}$ T cells were isolated and activated on anti-CD3, anti-CD28 and human-Fc-ICAM-1 coated plates. Cells were cultured in medium containing (Z)-4-Hydroxytamoxifen (4OHT). After 36 h, cells were harvested and treated or not with Bafilomycin A (BafA) for 2 h. Cell lysates had their ATG16L1 expression determined by immunoblotting. Data representative of 1 out of 4 experiments.

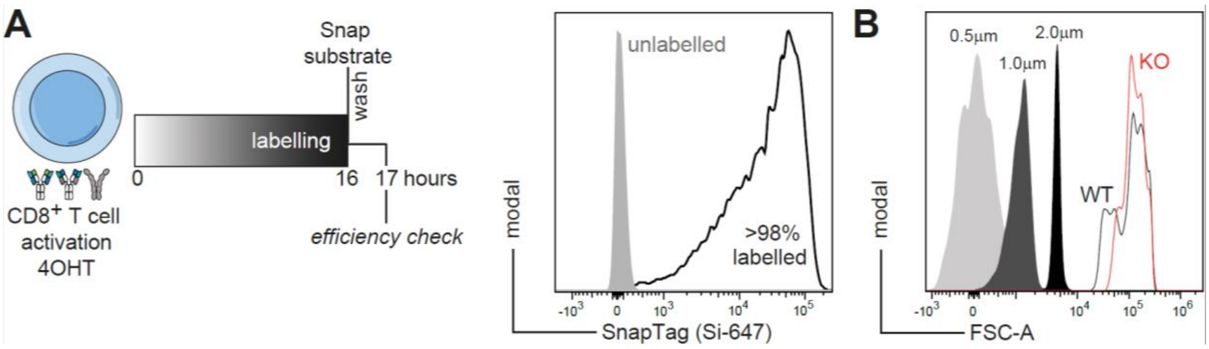


**Figure S2. SnapTag labelling is highly efficient and resistant to mitochondrial enrichment. (A)** Naïve MitoSnap CD8$^{+}$ T cells (*Omp25*-SnapTag$^{fl/-}$ *Ert2*$^{Cre}$) were cultured in medium containing (Z)-4-Hydroxytamoxifen (4OHT) and activated for 16 h prior to harvesting and labelling with a cell permeable SnapSubstrate (SNAP-Cell® 647-SiR). Efficiency of labelling was assessed by flow cytometry 30 min after substrate washing. **(B)** Representative histograms exhibiting beads of known size (0.5 µm, 1µm and 2µm) and mitochondrial populations enriched from *Omp25*-SnapTag$^{fl/-}$ *Ert2*$^{Cre}$ (WT) and *Omp25*-SnapTag$^{fl/-}$ *Atg16l1*$^{fl/fl}$ *Ert2*$^{Cre}$ (KO) cells. Cells were activated for 40 h and labelled with two SnapSubstrates (old and young mitochondria) as represented in Fig. 2A.

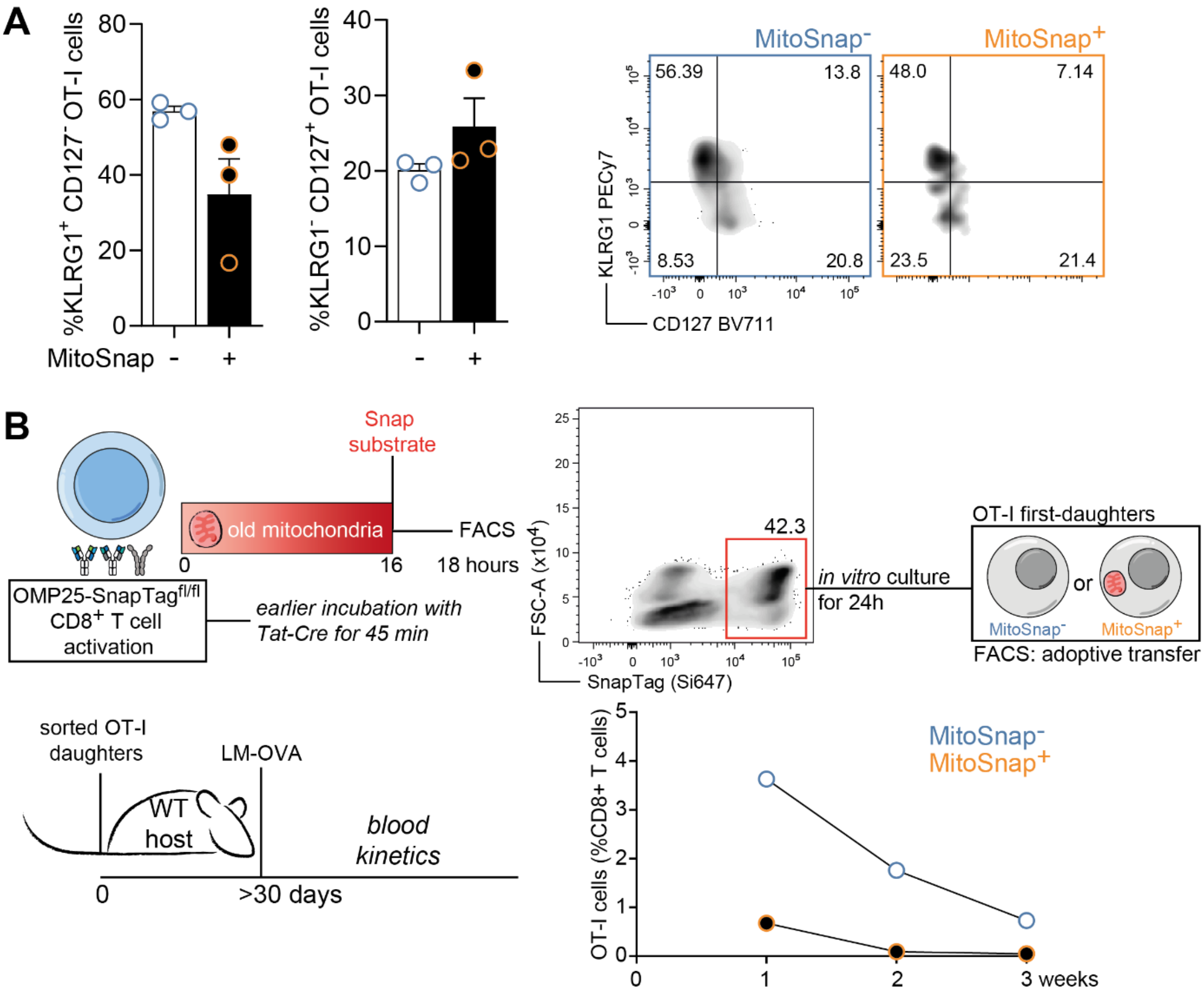


**Figure S3. MitoSnap+ and MitoSnap- progenies do not differ phenotypically at the memory-phase following cognate-antigen challenge, but the Tat-Cre recombinase system replicates results obtained using *Ert2*$^{Cre}$ concerning survival and re-expansion potential of these populations. (A)** OT-I MitoSnap CD8$^{+}$ T cells (*Omp25*-SnapTag$^{fl/fl}$ *Ert2*$^{Cre}$) were activated, labelled for old and young mitochondria (Fig. 2A) and sorted into MitoSnap+ and MitoSnap- prior to adoptive transfer ($5\times10^{4}$ cells intravenously) into new hosts. Progenies emerging from MitoSnap+ and MitoSnap- cells were monitored over the course of an immune response against *Listeria monocytogenes* expressing OVA (LM-OVA) (Fig. 3A). At 30 days post-challenge, phenotype of OT-I cells was evaluated by the expression of KLRG1 and CD127. Frequencies of short-lived effector cells (KLRG1$^{+}$ CD127$^{-}$) and memory-committed (KLRG1$^{-}$CD127$^{+}$) were calculated. Gating strategy is depicted on the right. Representative data of 1 out of 4 experiments. Data are represented as mean ± SEM. **(B)** Tat-cre driven-recombination was used as an alternative for the 4OHT-driven recombination using *Ert2*$^{Cre}$. Recombination efficiency was evaluated by SnapTag labelling. Cells that did express *Omp25*-SnapTag up to 16 h post Tat-cre recombination were sorted and cultured for further 24h. First-daughter OT-I MitoSnap+ and MitoSnap- cells were sorted and used in adoptive transfer experiments ($2\times10^{4}$ cells intravenously) (similar to Fig. 3A). Progenies emerging from MitoSnap+ and MitoSnap- cells were monitored over the course of an immune response against *Listeria monocytogenes* expressing ovalbumin (OVA) (LM-OVA). Representative data of 1 out of 2 experiments.

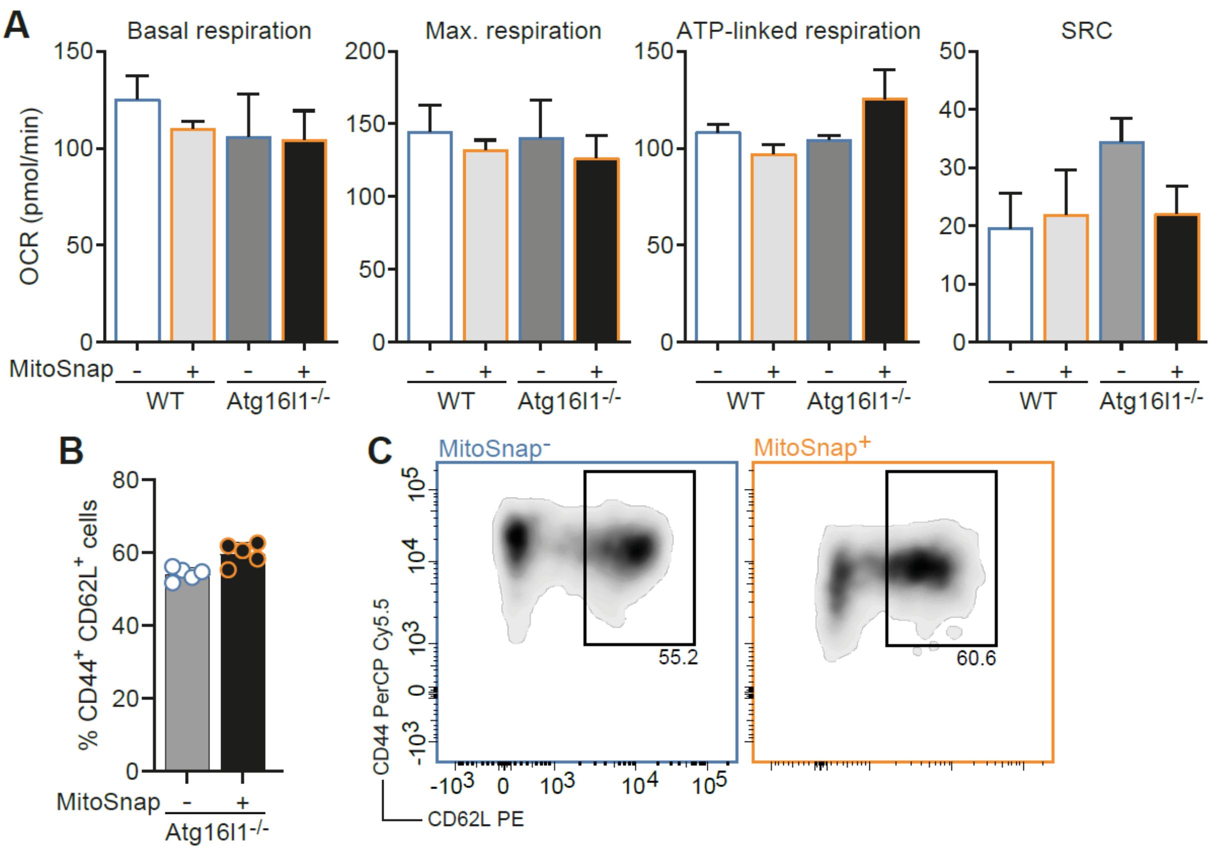


**Figure S4. MitoSnap+ and MitoSnap- progenies show similar oxygen consumption rate and autophagy-deficient daughter-cells do no differ phenotypically in survival assays. (A)** OT-I MitoSnap CD8+ T cells (*Omp25*-SnapTag$^{fl/fl}$ *Ert2*$^{Cre}$ or *Omp25*-SnapTag$^{fl/-}$ *Atg16l1*$^{fl/fl}$ *Ert2*$^{Cre}$) were activated, labelled for old and young mitochondria (Fig. 2A) and sorted into MitoSnap+ and MitoSnap- cells. Their oxygen consumption rate (OCR) was then measured using a XF96 MitoStress Test. Basal respiration, maximal respiration, ATP-linked respiration and spare respiratory capacity (SRC) were calculated. Data are represented as mean ± SEM. **(B)** MitoSnap CD8+ T cells (*Omp25*-SnapTag$^{fl/-}$ *Atg16l1*$^{fl/fl}$ *Ert2*$^{Cre}$) were activated, labelled for old and young mitochondria (Fig. 2A), sorted into MitoSnap+ and MitoSnap- populations after 36-40h and cultured for further 7 days in medium containing IL-2, IL-7 and IL-15. Surviving cells had their phenotype evaluated concerning the co-expression of CD44 and CD62L. Gating strategy is depicted on the right. Data are represented as mean ± SEM. Datapoints represent 5 technical replicates from 1 biological sample per group. Representative data from 1 out of 2 experiments.

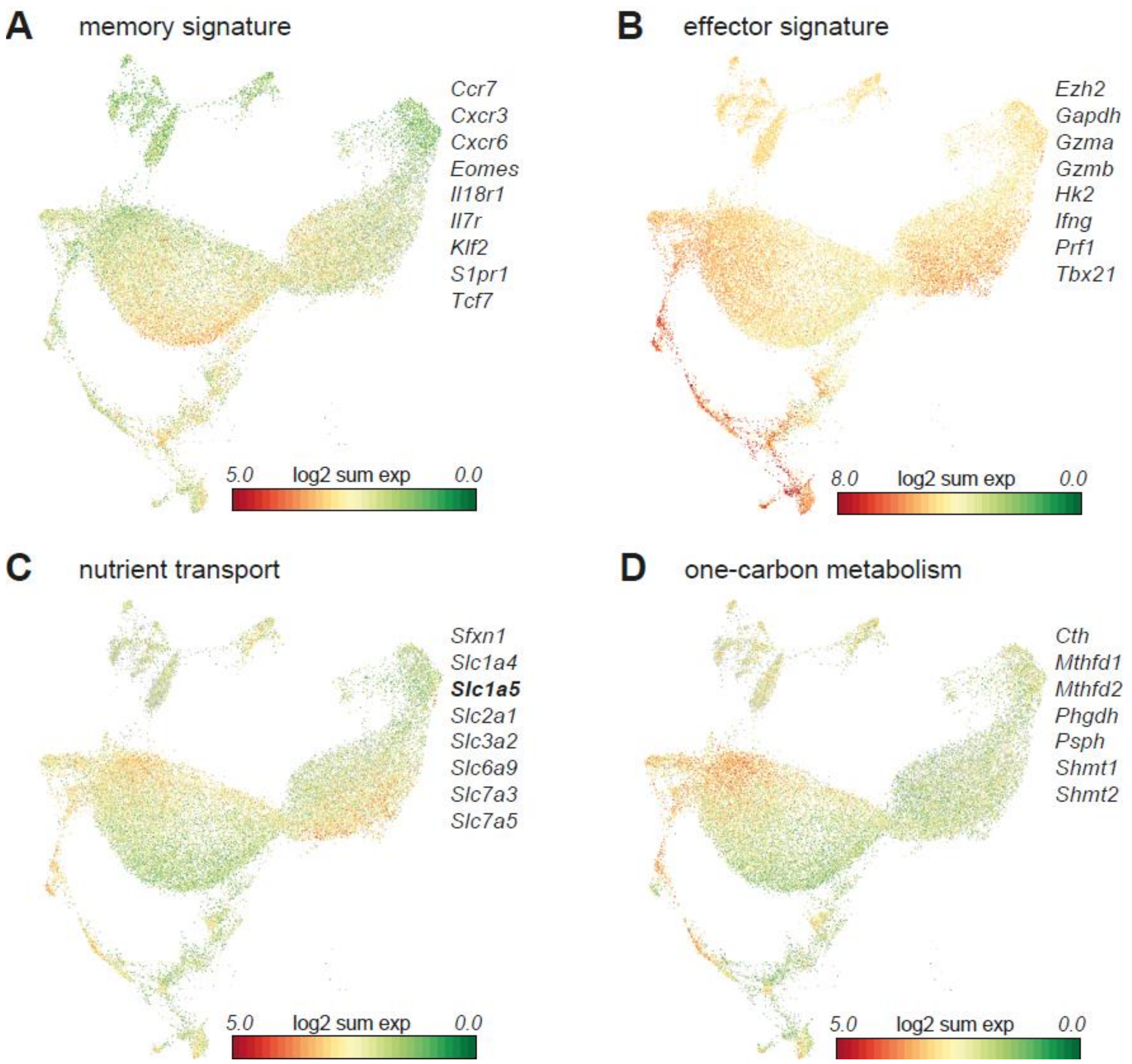


**Figure S5. MitoSnap+ and MitoSnap- progenies show distinct transcriptional profile.** OT-I MitoSnap CD8$^{+}$ T cells (*Omp25*-SnapTag$^{fl/fl}$ *Ert2*$^{Cre}$) were activated, labelled for old and young mitochondria (Fig. 2A) and sorted into MitoSnap+ and MitoSnap- populations. Single-cell transcriptomics analysis revealed that genes linked to **(A)** memory-fate commitment, **(B)** effector functions, **(C)** transport of glucose and amino acids and **(D)** involved in one-carbon metabolism are preferentially found in clusters enriched in MitoSnap+ or MitoSnap- cells (refer to Fig. 5 C). UMAP projections were extracted from Loupe Cell Browser.
